# Global translational and metabolic remodelling during iron deprivation in *Toxoplasma gondii*

**DOI:** 10.1101/2025.08.11.669662

**Authors:** Jack C. Hanna, Shikha Shikha, Megan A. Sloan, Clare R. Harding

## Abstract

Iron is required to support essential cellular processes. Due to diverse and dynamic host environments, the obligate intracellular parasite *Toxoplasma gondii* must adapt to iron limited conditions. To investigate the adaptations critical to parasite survival under these conditions, we conducted proteomic and metabolomic profiling of *Toxoplasma* cultured in iron depleted conditions. We find that iron depletion results in remodelling of the parasite proteome and triggers swift translational repression. This occurs prior to downregulation of the iron-regulated translation factor ABCE1, indicating an upstream, ABCE1-independent mechanism. In the context of repressed translation, we also observe a significant rewiring of energy metabolism. Iron depleted *Toxoplasma* have altered mitochondrial morphology and a profound reduction in mitochondrial respiration. Untargeted metabolomic analysis revealed tricarboxylic acid cycle (TCA) cycle dysregulation, characterised by accumulation of citrate and fumarate, both substrates of iron-dependent TCA cycle enzymes, and accumulation of glycolytic intermediates. We found iron deprived parasites continue to take up glucose and maintain glycolytic output, comparable to iron replete conditions. Limiting glucose availability either in culture media or by genetic ablation of glucose uptake caused a significant increase in sensitivity to iron restriction. Conversely, limitation of mitochondrially metabolised glutamine improved parasite fitness in iron depleted conditions. Together, our results establish iron as a key regulator of parasite translation and metabolic flexibility and demonstrate carbon source availability as important in *Toxoplasma* adaptation to iron deprivation.

**Importance:** This study determines the effects of iron deprivation on the parasite *Toxoplasma gondii. Using* proteomics and metabolomics, we reveal iron as a novel regulator of both translation and energy metabolism in *Toxoplasma,* underpinning the importance of this nutrient for essential cellular processes. We find that iron depletion introduces a metabolic bottleneck whereby parasites are dependent on glucose as their major carbon source. By modulating the parasite’s metabolism by altering carbon source availability, we identify nutrient conditions that improve parasite survival under iron restriction. These data reveal a key role for adaptive plasticity of *Toxoplasma* central carbon metabolism to drive survival under iron limited conditions. Understanding the interactions between parasite nutrient availability and metabolism allows us both to map the metabolic flexibility of these parasites, and to identify potential vulnerabilities.

## Introduction

Affecting approximately 1/3 of the global population, *Toxoplasma gondii* poses a persistent threat to human and animal health worldwide (Daher *et al*., 2021). A member of the apicomplexan phylum, *Toxoplasma* is related to other parasites of significant human and veterinary importance including *Plasmodium* (causative agent of malaria) and *Cryptosporidium* (causative agent of severe diarrheal disease). As an obligate intracellular parasite, *T. gondii* acquires essential nutrients from its host cell. These scavenged nutrients include carbon sources (e.g. glucose and glutamine), lipids, amino acids and vitamin cofactors (MacRae *et al*., 2012; Nolan, Romano and Coppens, 2017; Blume and Seeber, 2018; Fairweather *et al*., 2021). Unlike other members of its phylum, *T. gondii* demonstrates an unusually broad host cell range, able to infect all warm-blooded species and any nucleated cell type. This promiscuity is key to the success of the parasite, however the range of potential host environments necessitates flexibility in nutrient uptake, utilisation and in parasite metabolism (Blume *et al*., 2009; Nitzsche *et al*., 2016; Rajendran *et al*., 2021).

Iron is an essential nutrient for eukaryotic cells where it has been shown to be required for respiration, DNA replication, translation and metabolism (Evstatiev and Gasche, 2012), Underlining this importance, diverse cells have established mechanisms for surviving iron starvation, although with distinct regulatory processes. These mechanisms are often characterised by translational inhibition (Romero, Martínez-Pastor and Puig, 2021; Puig-Segui *et al*., 2024) and a metabolic switch away from oxidative phosphorylation towards glycolysis (Dziegala *et al*., 2018; Pereira *et al*., 2019; Chung *et al*., 2021). As mitochondrial respiration requires significant proportions of cellular iron stores (Jhurry *et al*., 2012), this metabolic switch allows cells to maintain energy generation under iron starvation. The proteomic remodelling and metabolic switch under iron depletion has significant functional consequences on cells, e.g. in macrophages, iron starvation-inducted metabolic changes reduce inflammation (Pereira *et al*., 2019).

Due to its central importance, mammals have developed mechanisms to starve pathogens of iron (Nairz and Weiss, 2020), making survival under iron depletion essential for successful pathogens. In *Toxoplasma*, iron is incorporated into a haem biosynthesis pathway (Bergmann *et al*., 2020) and three distinct iron-sulphur (Fe-S) cluster biogenesis pathways (Pamukcu *et al*., 2021; Renaud *et al*., 2024). These cofactors are essential components of proteins of mitochondrial metabolism and of the electron transport chain (ETC). Our previous work has identified widespread transcriptional changes upon acute iron deprivation (Sloan *et al*., 2025), however, changes in transcript abundance do not always correlate with functional adaptations, especially under stress (Liu, Beyer and Aebersold, 2016), which concealed any potential parasite adaptation to low iron.

To investigate the functional response of *Toxoplasma*, we combined global proteomics and metabolomics to assess how the parasite responded to acute iron depletion. We find iron depletion leads to rapid translational repression, accompanied by extensive proteomic remodelling. We assess the kinetics of repression and find translational repression precedes a drop in the abundance of ABCE1, a key iron-regulated component of effective translation. The proteomic changes were accompanied by a collapse in mitochondrial oxidative phosphorylation and significant metabolic changes in the cytosol and mitochondrion. Interestingly, we find that carbon source availability modulates the parasite’s sensitivity to iron deprivation; inhibition of glycolysis led to increased sensitivity, while glutamine restriction-which likely promotes energy generation through glycolysis-leads to increased growth under low iron. Our results demonstrate *Toxoplasma* actively responds to iron deprivation by remodelling its proteome, shifting energy generation to glycolysis, demonstrating the flexibility and adaptability of these intracellular parasites to nutrient stress.

## Results

### Iron deprivation leads to remodelling of the proteome

To investigate the impact of acute iron deprivation on the *Toxoplasma* proteome, we cultured *Toxoplasma* for 24 h in the presence of the iron chelator deferoxamine (DFO) on previously iron-starved host cells (pretreated for 24 h DFO) and quantified changes in parasite protein abundance by quantitative mass spectrometry (**Fig. 1A**). 5052 proteins were detected in at least two replicates and PCA analysis showed good grouping of samples between biological experiments, with clear separation between treatment conditions (**Fig. 1B**). Iron deprivation led to significant remodelling of the parasite proteome (**Fig. 1B, Table S1**), with 194 proteins significantly upregulated (adjusted *p* < 0.05, log_2_ fold change (LFC) > 0.6) and 365 significantly downregulated (adj. *p* < 0.05, LFC < -0.6).

**Figure 1.**
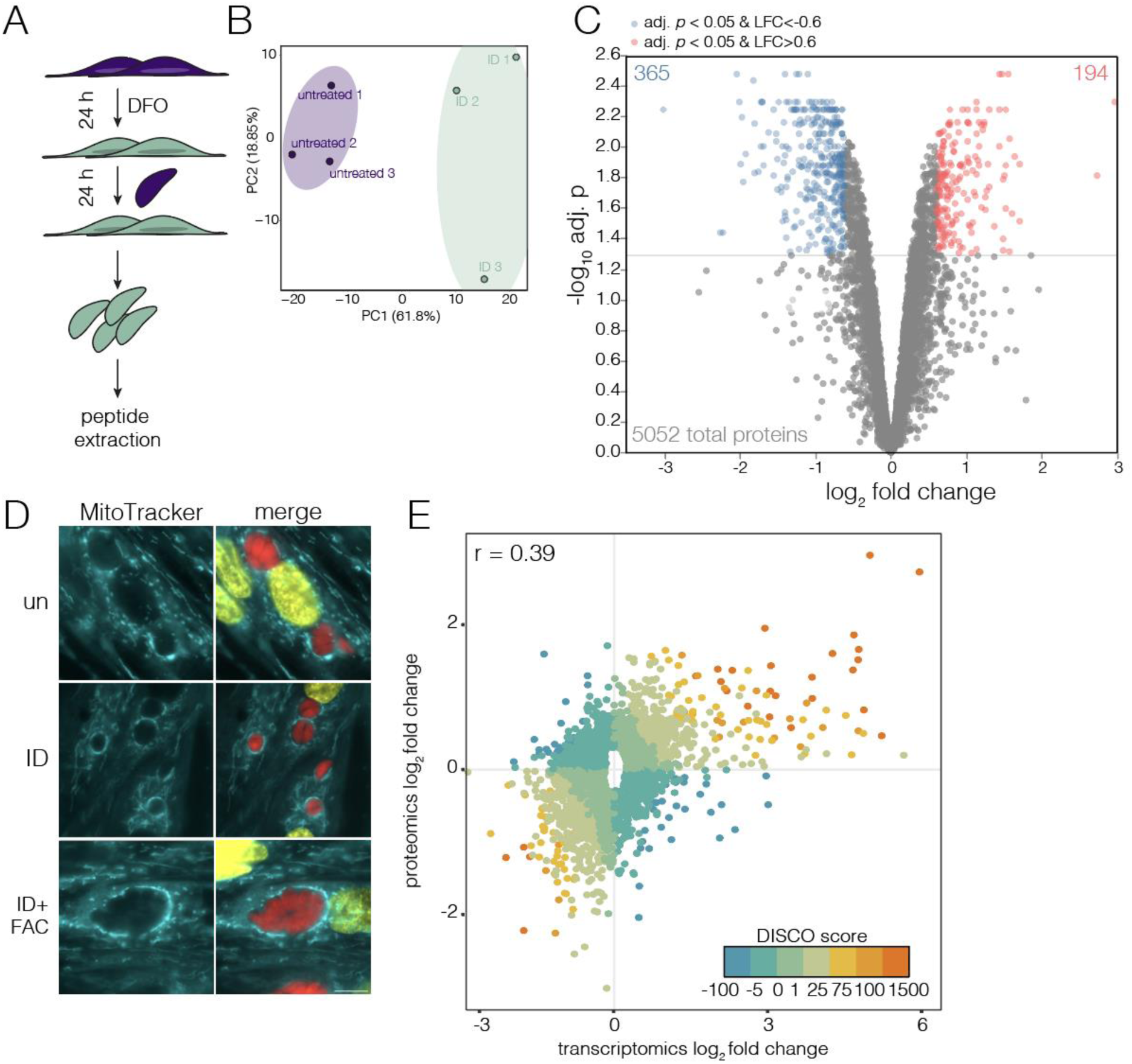
Iron deprivation leads to remodelling of the proteome. (**A**) Schematic showing sample collection workflow. Parasites were collected after 24 hours of infection on untreated host cells or cells that were pretreated with 100 µM DFO for 24 hours prior to infection (ID). (**B**) Principle components analysis (PCA) of untreated and iron depleted (ID) samples showing variation between biological replicates and conditions. Ellipses represent the 95% confidence region for each grouping. (**C**) Volcano plot of differential protein abundance analysis, with log_10_ adjusted *p*-value plotted against the log_2_ fold change (ID/untreated). Significant (adj. *p* <0.05, log_2_ fold change (L2FC) <-0.6 or >0.6) changing proteins indicated in red and blue. (**D**) Live imaging of host mitochondrial association (HMA) by MitoTracker in host cells cultured as above. For ID+FAC, host cells were pretreated with 100 µM DFO and 200 µM ferric ammonium citrate (FAC). MitoTracker staining localised adjacent to parasite vacuoles (red). Scale bar 10 µm. (**E**) Plot of proteomics log_2_ fold change against transcriptomics log_2_ fold change from (Sloan *et al*., 2025). Only genes that were significantly different in at least one dataset (adj.*p* < 0.05) were included in the analysis. A linear function was plotted (line not shown) to obtain correlation coefficient (r) of 0.39, points coloured by DISCO score. Genes which correlate positively between datasets have more positive (orange) DISCO scores, while those that are not correlated or negatively correlated have smaller or negative (blue) DISCO scores.

*Toxoplasma* encodes 70 predicted Fe-S-containing proteins (Renaud, Maupin and Besteiro, 2025) which gain Fe-S clusters through cytosolic, apicoplast or mitochondria-localised pathways (Pamukcu *et al*., 2021; Renaud *et al*., 2022). Disruption of Fe-S cluster biosynthesis pathways can result in destabilisation of the dependent Fe-S proteins (Pamukcu *et al*., 2021; Renaud *et al*., 2022; Maclean *et al*., 2024). We detect 66 of these predicted Fe-S proteins in our dataset, with 10 significantly changing in abundance in iron deprived parasites (**Fig. S1A**). Interestingly, we did not see depletion of proteins which gain Fe-S clusters from the mitochondrial or apicoplast pathways, however of the 8 Fe-S proteins known to require the cytosolic Fe-S pathway (Maclean *et al*., 2024), 5 were depleted in our dataset (**Table 1**). These data suggest that cytosolic Fe-S proteins may be more labile upon iron deprivation than organellar-localised proteins.

**Table 1.**
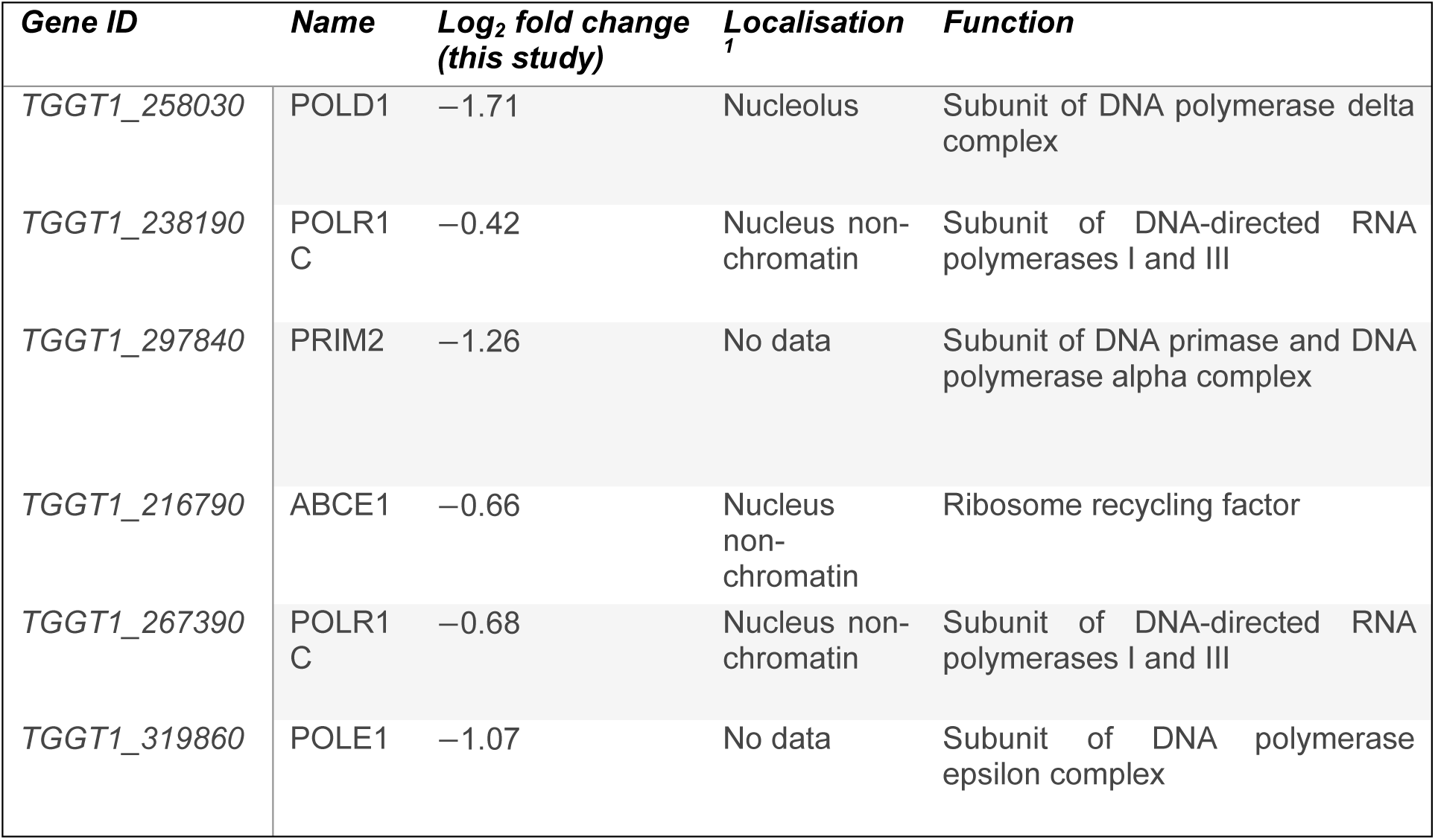
Cytosolic and nuclear Fe-S proteins (identified in Renaud, Maupin and Besteiro, 2025) which are downregulated in low iron. ^1^(Barylyuk et al., 2020)

Previously, iron starvation in *Toxoplasma* was shown to lead to an increase in lipid bodies (Renaud *et al*., 2024) and we see upregulation of the 2-acylglycerol O-acyltransferase (TGGT1_226370) that plays a crucial role in the formation of triglycerides that are stored within lipid droplets. *Toxoplasma* encodes pathways for both synthesis and scavenging of lipids. In our data, we see decreased abundance of ATP Citrate lyase (ACL - TGGT1_223840), used to synthesise acetyl-CoA from TCA-derived citrate, and a concurrent upregulation of acetyl CoA synthetase (ACS - TGGT1_266640). ACS was previously shown to generate cytosolic acetyl CoA from host derived acetate for fatty acid elongation (Dubois *et al*., 2018). Previous quantification of lipid species also found a decrease in sphingomyelin abundance (Renaud *et al*., 2024). However, we observed downregulation of both the serine palmitoyltransferase SPT1 (TGGT1_290980) and the ceramide synthase CERS2 (TGGT1_316450), both required for the *de novo* synthesis of ceramides and sphingomyelin. Previous data has shown that parasites preferentially scavenge sphingomyelins under nutrient replete conditions (Nyonda *et al*., 2022). An increase in sphingomyelin, despite the downregulation of synthesis enzymes, may therefore reflect an increase in sphingolipid scavenging by the parasite.

Taken together, these data indicate an increase in scavenging and decrease in synthesis of fatty acid building blocks, leading to increased storage of fatty acids within lipid droplets.

We did not detect changes in abundance of the major iron and zinc-uptake transporter (ZFT) or in the vacuolar iron transporter (VIT) (Aghabi *et al*., 2023, 2025), however the mitochondrial iron transporter (MIT – TGGT1_277090) did increase in abundance. Of the 194 proteins significantly upregulated, 70 of these were predicted to localise to the dense granules (DG). These proteins are typically secreted from the parasite to engineer a permissive environment for parasite growth (Griffith, Pearce and Heaslip, 2022). One potential hypothesis for accumulation of DG proteins in the parasite is reduced secretion under iron deprivation. To determine if dense granule secretion occurred in iron deprived cells, we examined host mitochondrial attachment (HMA) to the parasitophorous vacuole (PV). HMA is driven by the secretion of MAF1, a dense granule protein which specifically recruits host mitochondria (Pernas *et al*., 2014). However, even in iron deprived parasites we saw strong association of the host mitochondria to the PV (**Fig. 1D**). We also examined localisation of the DG protein Gra7 (Jacobs *et al*., 1998) and observed robust Gra7 secretion to the PV in iron deprived parasites (**Fig. S1B**), demonstrating that iron deprived parasites are capable of secreting DGs into the host cell.

Previously, we performed RNAseq on iron-deprived parasites under identical conditions (Sloan *et al*., 2025). To determine how changes in transcript abundance correlate with changes in the proteome, we correlated the fold changes in transcript or protein abundance after 24 h of iron deprivation (**Fig. 1E, Table S2)**. Points were coloured by discordance/concordance score (DISCO) based on the local FDR-corrected q-value for each L2FC (Domaszewska *et al*., 2017). We find that changes in protein abundance show limited correlation (Pearson’s correlation *r* = 0.39) with RNAseq results (**Fig. 1E**), and 1060 genes (34% of significantly changed genes) had a DISCO score of <0, suggesting high discordance between the approaches. Previous studies across organisms have seen disconnects between the transcriptome and proteome, especially under stress (Cheng *et al*., 2016; Teh *et al*., 2025) however in *Toxoplasma* this has not been examined in detail. Interestingly, in our transcriptomic data we see a general upregulation of transcripts, while at the proteome level we see overall downregulation of protein abundance. This discordance could be created by changes in translational rates under iron deprivation.

### Iron depletion leads to translational repression

Of the 365 proteins downregulated, 47 are involved in translation (**Fig. 2A**), and translation is a highly enriched GO terms among proteins with decreased abundance (**Fig. 2B**). To determine the functional consequence, we quantified total protein translation under iron deprivation using puromycin incorporation (Maclean et al., 2024). Puromycin is an aminonucleoside which is readily incorporated into nascent peptides during translation, allowing detection of puromycilated peptides with antibodies. After 24 hours of iron deprivation, we saw a significant (*p* value = 0.0015, one way ANOVA) decrease in global puromycin incorporation in the parasites after 30 mins of treatment, when normalised to total protein (**Fig. 2C and D**), demonstrating an inhibition of translation. To confirm this inhibition was due specifically to iron depletion, addition of exogenous iron (200 µM ferric ammonium citrate (FAC)) along with DFO led to no change in translation (**Fig. 2D**).

**Figure 2.**
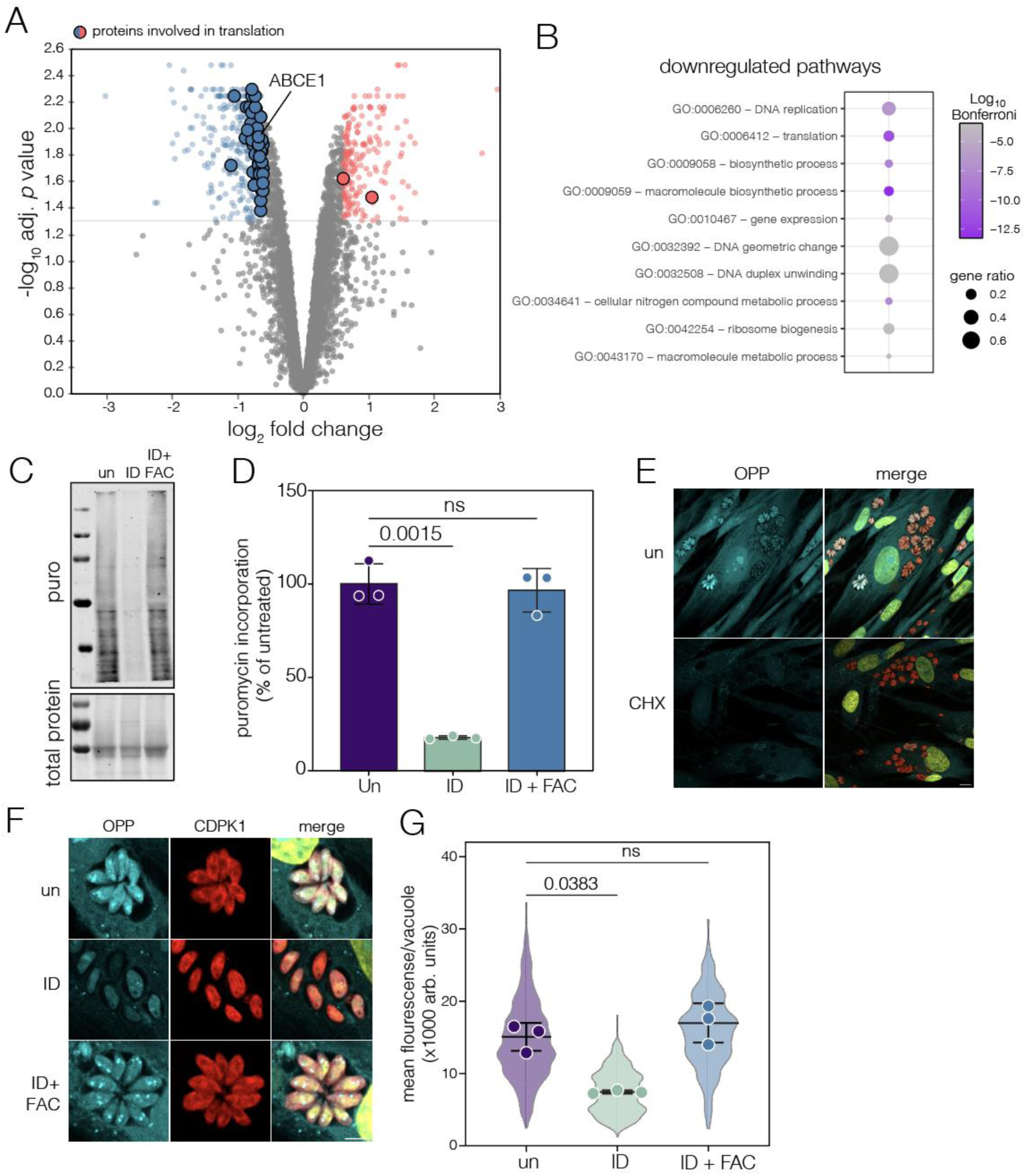
Iron depleted *Toxoplasma* downregulate translation. (**A**) Volcano plot from **Fig. 1C** with proteins significantly (adj. *p* < 0.05, L2FC <0.6) changed upon ID and involved in translation (using GO-term (GO:0006412)) highlighted. (**B**) Dot plot of the top 10 statistically significant GO terms from downregulated proteins, size represents of the proportion of genes annotated with each GO term. The colour intensity represents the Bonferroni corrected *p*-value. (**C**) Parasites, treated as indicated and incubated with puromycin for 30 minutes. Total protein visualised by ponceau S . Puromycilated peptides detected using an anti-puromycin antibody. (**D**) Quantification of puromycin incorporation, intensity normalised to total protein and untreated mean. Each point represents a biological replicate, bar at mean±SD *p* values from Brown-Forsythe and Welch ANOVA. (**E**) Immunofluorescence images of *Toxoplasma* vacuoles labelled with o-propargyl puromycin (OPP) (cyan), with and without co-incubation of cycloheximide (CHX). CDPK1 used as parasite cytoplasmic control. Scale bar 10 µm. (**F**) Immunofluorescence imaging of OPP in untreated (top) 24 hour ID (middle) and ID+FAC (bottom) *Toxoplasma*. Scale bar 5 µm. (**G**) Quantification of OPP incorporation/vacuole. Each point represents a biological replicate, bar at mean±SD, *p* values from one way ANOVA with Tukey’s multiple comparison correction. Violins represent the distribution of all vacuoles measured from each condition (N=537, 573 and 476 respectively).

Puromycin will also be incorporated by the host cells, which will also be impacted by iron deprivation (Puig-Segui *et al*., 2024), potentially complicating our results. To validate the effect on the parasites, we used an analogue of puromycin, O-propargyl puromycin (OPP) (Liu *et al*., 2012) which allows for the covalent attachment of fluorophores to puromycilated peptides by click chemistry. Using an attached fluorophore, we were able to visualise OPP-incorporation (**Fig. 2E**). OPP-incorporation was seen in both the host and parasites, and was heterogeneous between vacuoles, potentially reflecting the asynchronous cell cycle of *Toxoplasma* vacuoles (Batra *et al*., 2024). To confirm specificity of OPP, incorporation could be effectively blocked by cycloheximide (CHX) (**Fig. 2E, Fig. S2A**), an inhibitor of cytosolic translation. Quantifying OPP incorporation demonstrated significantly (*p* = 0.038, one way ANOVA) reduced signal in iron deprived vacuoles, which was not seen upon DFO and iron co-treatment (**Fig. 2F and G**).

These data confirm that iron deprivation induces global translational repression in *Toxoplasma*.

### Translational repression is rapid and precedes ABCE1 depletion

Of the Fe-S proteins found to be significantly downregulated upon iron deprivation (**Table 1**), ABCE1 has a well-documented role in ribosome recycling during translation (Sudmant *et al*., 2018) and is iron-regulated in mammalian cells (Sudmant *et al*., 2018; Zhu, Zhang and Mendell, 2020). There is conflicting data on the result of depletion of ABCE1 on translation in mammalian cells, with some studies showing this is sufficient to repress translation (Toompuu *et al*., 2016; Annibaldis *et al*., 2020) while others see only a minor effect (Zhu, Zhang and Mendell, 2020).

In our dataset, ABCE1 is significantly depleted (**Fig. S2B**) (Log_2_FC = -0.66, FDR-adjusted *p =* 0.012). To confirm this, we used an endogenously ABCE1-HA-tagged line (Maclean *et al*., 2024) and saw ABCE1-HA was significantly decreased (*p* = 0.0118, one way ANOVA) under low iron by western blot (**Fig. 3A and B**), normalised to CDPK1, a highly abundant protein unchanged in proteomics (**Table S1**). We confirmed by immunofluorescence assay and saw a significant and reproducible (*p* = 0.0019, one way ANOVA) drop in ABCE1-HA signal upon iron deprivation (**Fig. 3C and D**).

**Figure 3.**
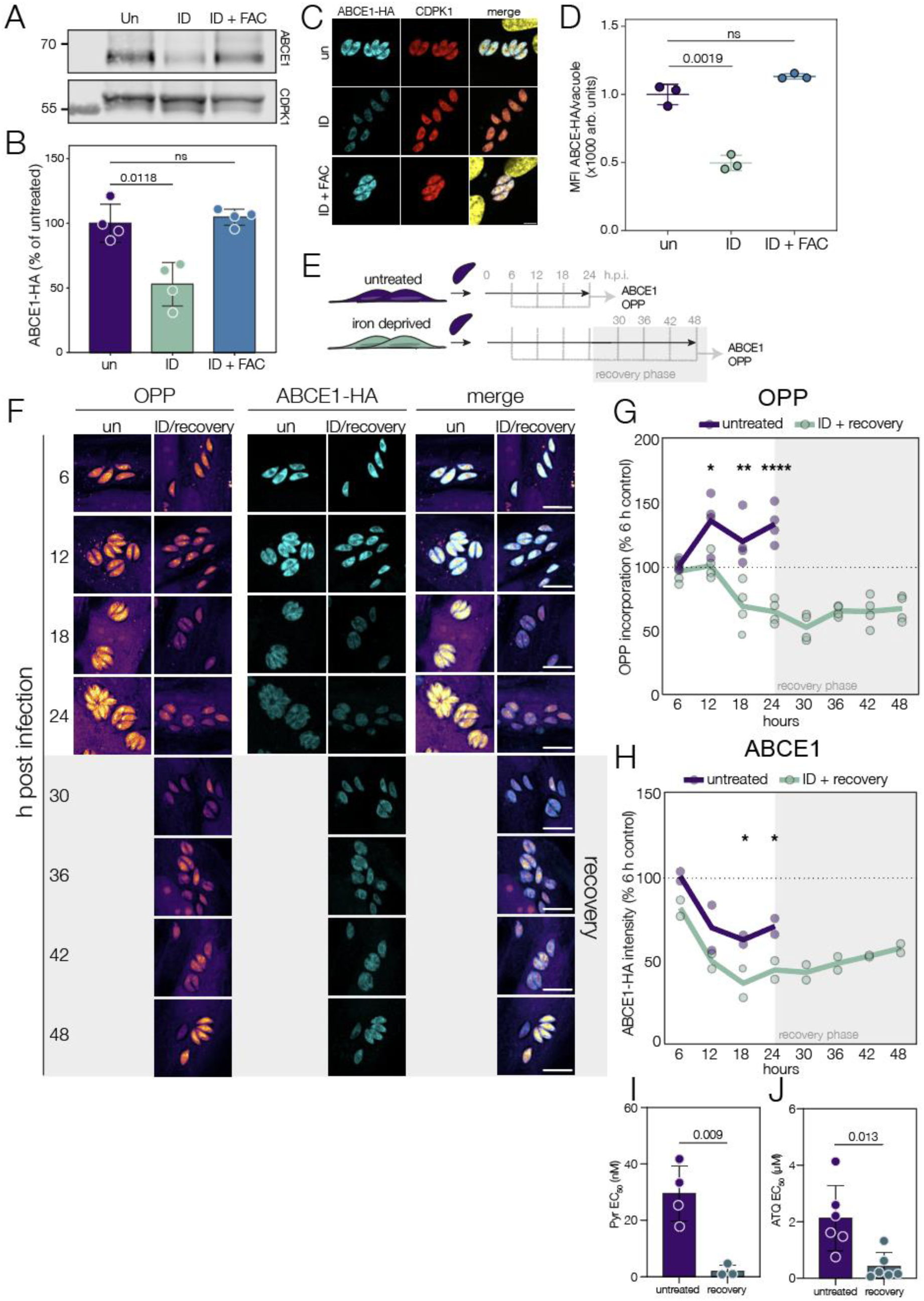
Translational repression is rapid, and precedes ABCE1 depletion. (**A**) Immunoblot of ABCE1-HA lysates from untreated, ID or ID+FAC parasites. CDPK1 included as a loading control. (**B**) ABCE1-HA abundance was quantified and normalised to CDPK1 abundance and untreated parasites, points indicate biological replicates, bars at mean±SD, *p* value from one-way ANOVA with Dunnet’s correction for multiple comparisons. (**C**) Immunofluorescence of ABCE1-HA parasites grown as above, CDPK1 included as cytoplasmic marker. ABCE1-HA localises to cytoplasm. Scale bar 5 µm. (**D**) ABCE1-HA signal quantified from 3 biological replicates, bars at mean±SD. *p* value from Brown-Forsythe and Welch ANOVA. (**E**) Schematic depicting the translation imaging time course. Samples from untreated and ID parasites were imaged every 6 hours for 24 hours. Additional samples of ID parasites which then cultured in iron replete media for a further 24 hours up to 48 hours post infection included to assess translation recovery. (**F**) Immunofluorescence images of ABCE1-HA parasites every 6 hours through ID and recovery. OPP incorporation (left) and ABCE1 (middle) were quantified. Scale bar 10 µm. Mean (lines) of OPP (**G**) and ABCE1-HA signal (**H**) plotted relative to untreated 6h. *p* values from comparing ID with untreated, two way ANOVA with Tukey correction, * *p* < 0.03, ** *p* < 0.003, *** *p* < 0.0001 (**I**) Average EC_50_ of pyrimethamine and atovaquone (**J**) against untreated and parasites recovering from iron depletion. Points indicate biological replicates, bars at mean ±SD. *p* value from two tailed t test.

To examine the kinetics of ABCE1 depletion in the context of translational repression, we performed a time course of iron deprivation and subsequent recovery when parasites are given iron replete media after 24h of starvation (**Fig. 3E**). At each timepoint, we quantified vacuolar area, OPP incorporation (as a measure of translation) and ABCE1-HA abundance in each vacuole. Mean vacuolar area did not increase during iron depletion, as expected (**Fig. S2C**). However, replacement with iron replete media after 24 h iron depletion did lead to a significant (*p* = 0.03, one way ANOVA) increase in vacuolar area from 12 h post recovery, indicating that parasites were able to reinitiate replication upon recovery, and demonstrating the effects of acute iron depletion are largely reversible.

We also quantified host cell OPP-incorporation throughout the experiment. Host translation was notably lower than parasites at all points, and iron depletion significantly reduced host translation (*p* < 0.0001, two way ANOVA with Tukey’s correction) at each time point, as previously described (Puig-Segui *et al*., 2024). Host translation recovered by 36 h post treatment (12 h post recovery) with no significant change from untreated host cells (**Fig. S2D**)

In the parasite, levels of OPP incorporation are consistent between vacuoles at early time points, but become more diverse after 6 h, potentially reflecting the progressive loss of synchronous cell division (**Fig. S2E**). Upon iron depletion, the first significant (*p* = 0.039, two way ANOVA with Tukey’s correction) difference in translation appears at 12 hours post infection (**Fig. 3F, G**), as iron deprived parasites no longer increase translation. After 12 h of iron depletion, levels of OPP incorporation progressively decrease (**Fig. 3F, G**). However, upon recovery, mean OPP-incorporation significantly (*p* = 0.034, two way ANOVA with Tukey’s correction) increases (**Fig. S2F**), demonstrating recovery of translation under iron replete conditions.

Recent single cell transcriptomics identified ABCE1 as cell cycle regulated, with a wave of transcription occurring in G1-S phase (Lou *et al*., 2024). We see a peak of ABCE1-HA abundance at 6 h post infection, which then drops in the untreated cells (**Fig. 3F, H**). There is no significant change in ABCE1-HA abundance under iron depletion until 18 hours (*p* = 0.03, two way ANOVA with Tukey’s correction), despite the drop in translation. Upon recovery, there is a gradual (although not significant) increase in ABCE1 abundance over the course of recovery (**Fig. S2F**).

The stall in translation, prior to ABCE1 abundance changes, and the failure of iron replete parasites to recover ABCE1 levels, suggests that loss of ABCE1 is not the major driver of translational repression in iron depleted parasites.

Addition of iron to deprived parasites triggered a recovery of translation and replication. To determine if this recovery led to an increased reliance on key metabolic pathways, we iron deprived parasites for 24 h, replaced with iron replete media and tested sensitivity to metabolic inhibition. Recovering parasites were significantly (*p* = 0.009, *t* test) more sensitive to pyrimethamine, an inhibitor of folate biosynthesis required for nucleotide synthesis (**Fig. 3I, S2G)**, and to inhibition of the mitochondrial ETC by atovaquone (*p* = 0.013, *t* test) (**Fig. 3J, S2H**). However, we saw no change in sensitivity upon treatment with the broad anti-parasitic dihydroartemisinin (**Fig. S2I, J**). This suggests that upon recovery from iron deprivation, parasites are more sensitive to further metabolic inhibition.

### Iron deprivation leads to alterations in mitochondrial metabolism

Beyond translation, iron is essential in the mitochondria where its incorporation into the proteins of the electron transport chain (ETC) is required for mitochondrial respiration and energy generation (Huet *et al*., 2018; Leonard *et al*., 2023; Silva *et al*., 2023). *Toxoplasma* has a single dynamic, mitochondrion which has been shown to change morphology based on stress (Vizcarra *et al*., 2023), genetic ablation of key components (Huet *et al*., 2018; Mallo *et al*., 2021) and upon extracellular exposure (Ovciarikova *et al*., 2017). We examined parasite mitochondrial morphology at 24 hours post iron starvation and saw that mitochondria from iron depleted parasites had lost the circular “lasso” shape observed in untreated parasites (**Fig. 4A**). This phenotype was fully reversible, as the mitochondrion reverted to normal after a further 24 hours in iron replete media (**Fig. 4A**). To understand the kinetics of mitochondrial morphology, we collected immunofluorescence images from untreated and iron deprived parasites every 6 hours for 24 hours of infection and 24 hours of recovery and assigned mitochondria into 3 previously established morphology classes (Ovciarikova *et al*., 2017) (**Fig 4B**). At both 6- and 12-hours post infection, mitochondrial morphology is unchanged. However, at 18 hours, iron depletion causes a significant decrease in the proportion of lasso mitochondria compared to untreated parasites (*p*=0.046 at 18 h, *p*=0.0022 at 24 h, two-way ANOVA) (**Fig. 4C**). Upon recovery, after 6 hours of culture in iron replete, there was a significant increase in the number of lasso shaped mitochondria (*p*=0.005, two-way ANOVA), reflective of the proportions seen in untreated intracellular parasites (**Fig.4D**).

**Figure 4.**
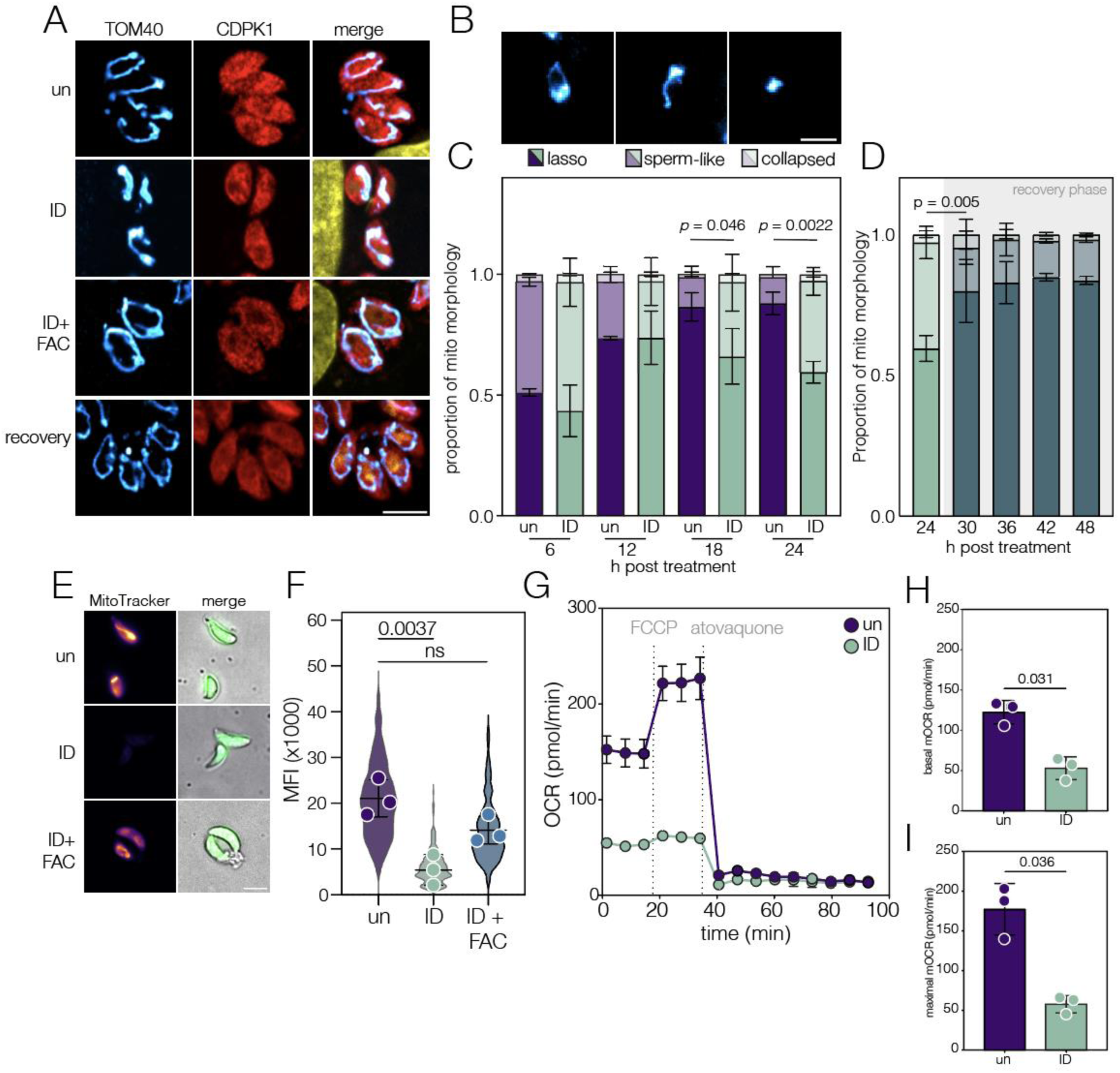
Iron deprivation leads to changes in mitochondrial metabolism. (**A**) Immunofluorescence of intracellular parasites treated as indicated. Mitochondrial morphology observed using the mitochondrial membrane marker TOM40, parasite cytoplasm labelled using anti-CDPK1. Scale bar 5 µm. (**B**) Mitochondrial morphologies, based on (Ovciarikova *et al*., 2017). Scale bar 5 µm. (**C**) Quantification of mitochondrial morphology through a control or ID infection, 100 parasites quantified in each replicate, N=2, *p* value from two way ANOVA with Tukey correction for multiple comparisons. (**D**) Stacked bar graph mitochondrial morphology across ID and recovery. Bars at mean±SD, *p* value from two way ANOVA with Tukey correction for multiple comparisons. (**E**) Representative live imaging of MitoTracker Deep Red staining in extracellular ΔKu80:mNeonGreen *Toxoplasma* grown in ID or ID+FAC conditions. Merged images show cytoplasmic mNeon and a polarised light image. Scale bar 5 µm (**F**) Quantification of Mitotracker intensity across 3 biological replicates (points), overlayed on violin plot showing the distribution of individual parasites measured (N=294/262/305). *p* values from one-way ANOVA with Tukey correction for multiple comparisons. (**G**) Representative oxygen consumption rate (OCR) trace for untreated and ID parasites. Background OCR was subtracted to determine mitochondrial associated oxygen consumption rate (mOCR). (**H**) Basal and (**I**) maximal mOCR was determined for both untreated and iron depleted parasites after 24 hours of infection. Each point represents a replicate (N=3), bar at mean ±SD, *p* values from two-tailed paired t tests.

As mitochondrial shape is linked to function, dynamic changes to morphology may indicate changes in function. We first assessed mitochondrial membrane potential using MitoTracker, a dye which accumulates in mitochondria and fluoresces in the presence of an active proton gradient. We observed a significant (*p* = 0.0037, one way ANOVA) decrease in MitoTracker signal in DFO-treated parasites (**Fig. 4E** and **F**).

To gain an integrated view of cellular energy metabolism, we performed metabolic flux analysis using a Seahorse Analyser (Hayward *et al*., 2022; Silva *et al*., 2023) which allows quantification of the mitochondrial oxygen consumption rate (mOCR). We find iron deprivation leads to a significant drop both in basal mOCR (**Fig. 4G and H**, *p* = 0.031, *t* test) and maximal mOCR (**Fig. 4I**, *p* = 0.036, *t* test), demonstrating a significant defect in mitochondrial respiratory capacity upon iron starvation.

These data demonstrate iron deprivation leads to significant defects in mitochondrial polarisation and respiration.

### Iron deprivation leads to global metabolic changes

Mitochondrial respiration is central to cellular metabolism, however many metabolic processes beyond the mitochondrion rely on iron-bound enzymes. To understand the global effects of iron depletion on parasite metabolism, we performed untargeted metabolomics on iron depleted parasites. 675 metabolites were annotated across four biological replicates, and the PCA plot (**Fig. 5A**) demonstrated good separation between experimental conditions. Overall, we identified 56 upregulated metabolites (adj. *p* < 0.1, log_2_ fold change (LFC) > 0.6) and 60 significantly downregulated (adj. *p* < 0.1, LFC < -0.6) (**Fig. 5B, C, Table S3**). Under iron deprivation, both deferoxamine and the iron-bound ferrioxamine were detectable; we find that approximately 65% of the chelator was iron bound (**Fig. 5D**), demonstrating that DFO is in excess under these conditions.

**Figure 5.**
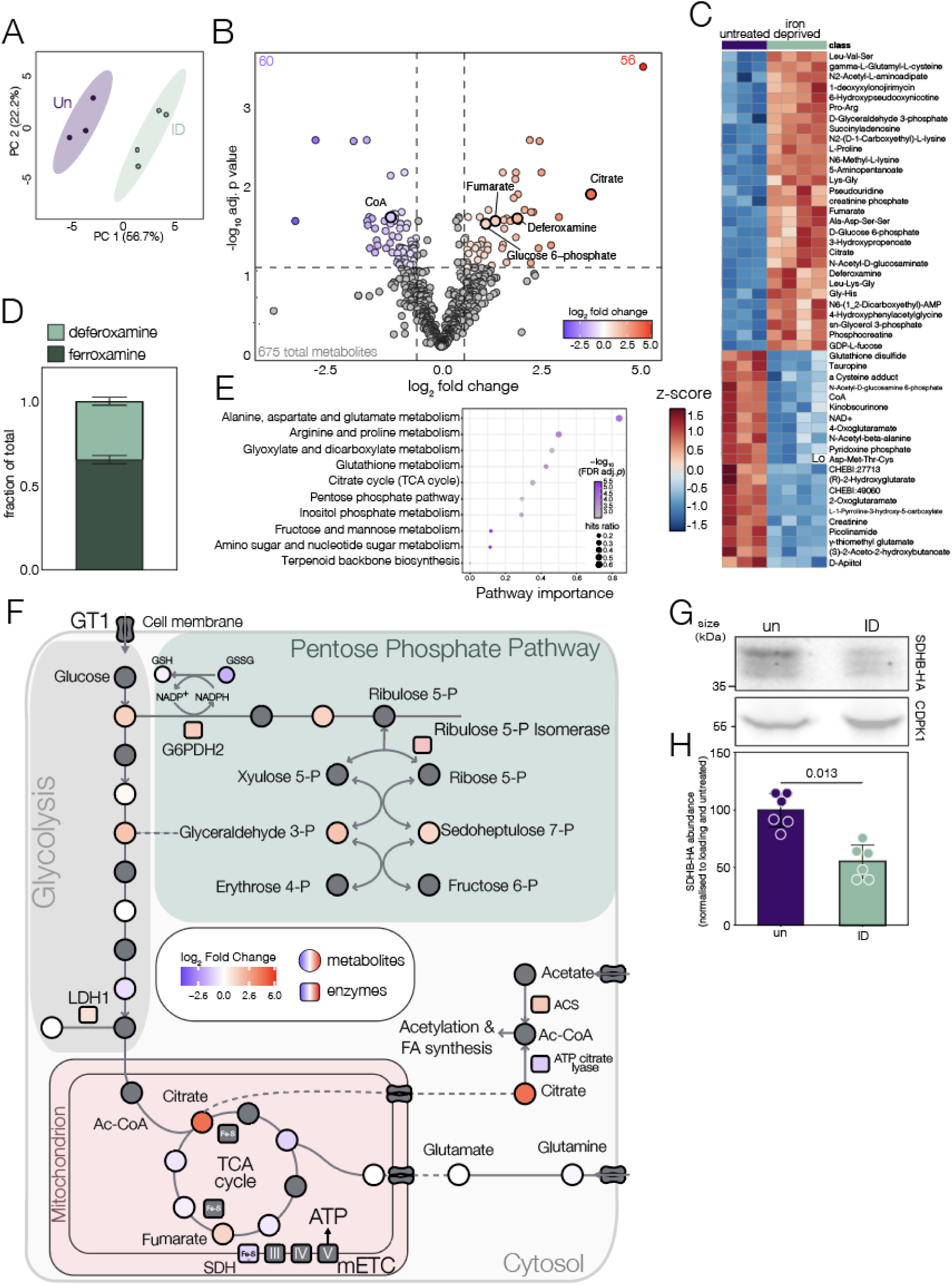
Iron deprivation leads to global metabolic changes. (**A**) PCA plot showing variation between replicates and untreated (purple) and iron depleted (green) groups. Ellipses represent the 95% confidence interval of each grouping. (**B**) Volcano plot showing identified metabolites log_10_ adjusted p-value (computed by FDR adjusted t-tests) against corresponding log_2_ fold change (ID/untreated). Metabolites with adjusted p<0.1 and a log_2_ fold change of >0.6 or <-0.6 are coloured based on magnitude of fold change, non-significant metabolites in grey. (**C**) Heatmap of 50 most significantly different metabolites, clustered on Euclidian distance, coloured by z-score transformed fold change. (**D**) Stacked bar graph showing the fraction of total iron chelator detected that is bound (ferroxamine) or unbound (deferoxamine) by iron. Normalised peak intensities for each divided by their combined totals for each replicate. (**E**) Dot plot of 10 most significantly enriched *Toxoplasma* KEGG pathways. Pathways are ranked by pathway importance, computed from pathway topology analysis, with pathways containing closely related metabolites assigned greater importance. Dot size represents the proportion of metabolites for each term represented among significant metabolites and colour represents -log_10_(FDR adjusted *p* value). (**F**) Schematic of central carbon metabolism with proteins represented by squares and metabolites represented by circles. Significantly different proteins are coloured, while all detected metabolites are coloured with the log_2_ fold changes metabolomics datasets. Metabolites not detected are represented by grey circles, white indicates detection but no significant difference. Asterisk (*) represents the fold change of SDHB determined by western blot. (**G**) and (**H**) immunoblot of lysates from SDHB-HA parasites, untreated or ID, CDPK1 included as loading control. Points represent biological replicates (N=6), bar at mean±SD, *p* value from two-tailed paired t test.

To identify the pathways of importance in our dataset, we performed pathway analysis of our identified metabolites. We incorporated pathway topology in our analysis so pathways with closely associated metabolites are assigned a higher “pathway importance” score (**Fig. 5E**). Important pathways identified by this analysis relate to several amino acid metabolic pathways including alanine, aspartate and glutamate metabolism along with arginine and proline metabolism. Iron depletion has previously been associated with perturbations to amino acid metabolism (Ying *et al*., 2024; Teh *et al*., 2025), however the cause of these changes in *Toxoplasma* is not yet understood. We see increased abundance of several di- and tri-peptides and an increase in post translationally modified N6-methyl lysine. Of the metabolites with the largest increases in abundance we identified 5-aminopentanoic acid and lysopine, both of which are lysine degradation derivatives (Knorr *et al*., 2018). which suggests that protein turnover imbalance brought about by translational repression may play a role.

One of the pathways significantly impacted by iron deprivation is the pentose phosphate pathway, which acts both to funnel carbon from glycolysis into nucleotide production and to reduce nicotinamide adenine dinucleotide phosphate (NADP) (**Fig. 5F**). We see increased abundance of several metabolites and proteins of the pathway, including glucose-6-phosphate dehydrogenase 2 (G6PDH2). Upregulation of G6PDH2 is protective against oxidative stress (Xia *et al*., 2022) and, via NADP recycling, is important to reduce glutathione disulfide (GSSG) to glutathione. Glutathione has been linked to signalling in iron starvation in multiple eukaryotes (Kumar *et al*., 2011; Shee *et al*., 2022), although its role in role in *Toxoplasma* is not yet fully understood.

We also see significant changes in fumarate and citrate, both metabolites of the TCA cycle, with a 14-fold increase in citrate abundance in iron depleted parasites. Citrate production in *Toxoplasma* occurs in the cytoplasm by ATP-citrate lyase (TGGT1_223840), which is significantly reduced in iron deprivation (L_2_FC -0.832, adj. *p* value = 0.0353), suggesting there is not an increase in citrate production. Citrate is isomerised iso-citrate by the Fe-S enzyme aconitase. Iron depletion is associated with changes in aconitase enzymatic function in mammalian systems (Castello, Hentze and Preiss, 2015; Cho *et al*., 2021) and we have shown inhibition of activity in iron deprived *Toxoplasma* (Sloan *et al*., 2025). Aconitase is essential for the parasite, however to test if the importance of aconitase enzyme activity changes under iron limitation, we determined the sensitivity of normal or iron deprived parasites to sodium fluoroacetate (NaFAc), a specific inhibitor of aconitase (MacRae *et al*., 2012; Harding *et al*., 2020). Due to the lack of replication at 100 µM DFO, we treated parasites with 20 µM DFO-predicted to inhibit parasite replication by around 20%-to limit iron but allow for parasite growth. However, 20 µM DFO-treated parasites become significantly (*p* < 0.0001, extra sum-of-squares F test) more sensitive to NaFAc (IC_50_ diff = -37.33 nM) (**Fig. S3A**), suggesting that while the TCA cycle is inhibited in activity, it remains important for parasite replication.

Fumarate is formed by the succinate dehydrogenase complex and is increased 2.6-fold in iron deprivation. To investigate this further, we examined abundance of key component, succinate dehydrogenase B (SDHB) (TGGT1_215280). Previously, we saw that knockdown of the major iron transporter ZFT lead to decreased SDHB abundance (Aghabi *et al*., 2025). Using a parasite line expressing endogenously tagged SDHB (SDHB-HA) (Maclean *et al*., 2021), we determined that SDHB was significantly (*p* = 0.013, *t* test) downregulated under iron depletion (**Fig. 5G and H**). This appeared specific, as a similar Fe-S protein of the ETC, Rieske-FLAG (MacLean *et al*., 2025) did not change in abundance under low iron conditions (**Fig. S3B and C)**, potentially indicating that these proteins differ in their stability under stress, or suggesting targeted regulation of SDHB. Despite the loss of SDHB, fumarate accumulates. Fumarate is subsequently hydrated to malate by fumarate hydratase. Unlike their mammalian hosts who utilise Fe-independent class II fumarate hydratases, *Toxoplasma* encodes a class I fumarate hydratase (TGGT1_267330) which does not change significantly in the proteomics. These class I fumarate hydratases contain a 4Fe-4S cluster present in prokaryotes and unicellular eukaryotes like typanasomatids and apicomplexans (de Pádua *et al*., 2017; Jayaraman *et al*., 2018; Feliciano, Drennan and Nonato, 2019) and have been shown to be inhibited by iron deprivation (Park and Gunsalus, 1995; Tseng, 1997). We suggest that in low iron, fumarate hydratase may lose activity (potentially due to Fe-S cluster loss), despite sustained protein abundance, leading to fumarate accumulation in low iron.

Iron deprivation leads to large changes in the parasite’s metabolic profile. Here, we link this with changes in metabolic enzyme abundance and activity. However, *Toxoplasma* is metabolically flexible and able to adapt metabolic pathways to changes in environmental conditions (Nitzsche *et al*., 2016; Dubois *et al*., 2018; Rimple *et al*., 2025).

### Glutamine restriction is protective under low iron

Metabolomics revealed significant changes to parasite central carbon metabolism under low iron conditions, however, does not reveal if distinct central carbon pathways become more fitness conferring under iron starvation. To determine their significance to parasite survival and replication, we assessed how altering parasite carbon sources changed sensitivity to iron deprivation. *Toxoplasma* takes up glutamine from their environment which supplies carbon to the TCA cycle (MacRae *et al*., 2012). As we saw changes in metabolites of the TCA cycle (**Fig. 5E**), we tested the sensitivity of glutamine-restricted (0.4 mM glutamine, 10% of normal conditions) parasites to iron depletion. *Toxoplasma* can grow in the absence of glutamine (Nitzsche *et al*., 2016), however we found removal of glutamine and limitation of iron caused reduced host cell fitness. However, restriction of glutamine led to a significant (Extra-sum-of-squares F test, *p* <0.0001) protective effect for iron deprived parasites, with significantly more DFO required to restrict parasite growth (IC_50_ diff = +20.75 µM) (**Fig. 6A and B**). We confirmed this result by plaque assay; although there was no significant effect on plaque number (**Fig. S4A**), limiting glutamine had no impact on parasite growth, while iron deprivation caused a significant decrease in plaque area, as expected (Nitzsche *et al*., 2016) (**Fig. 6C**). However, glutamine restriction was able to partially rescue (*p* = 0.018, one way ANOVA) plaque area upon iron deprivation (**Fig. 6C and D**), suggesting that glutamine restriction is protective for iron deprivation.

**Figure 6.**
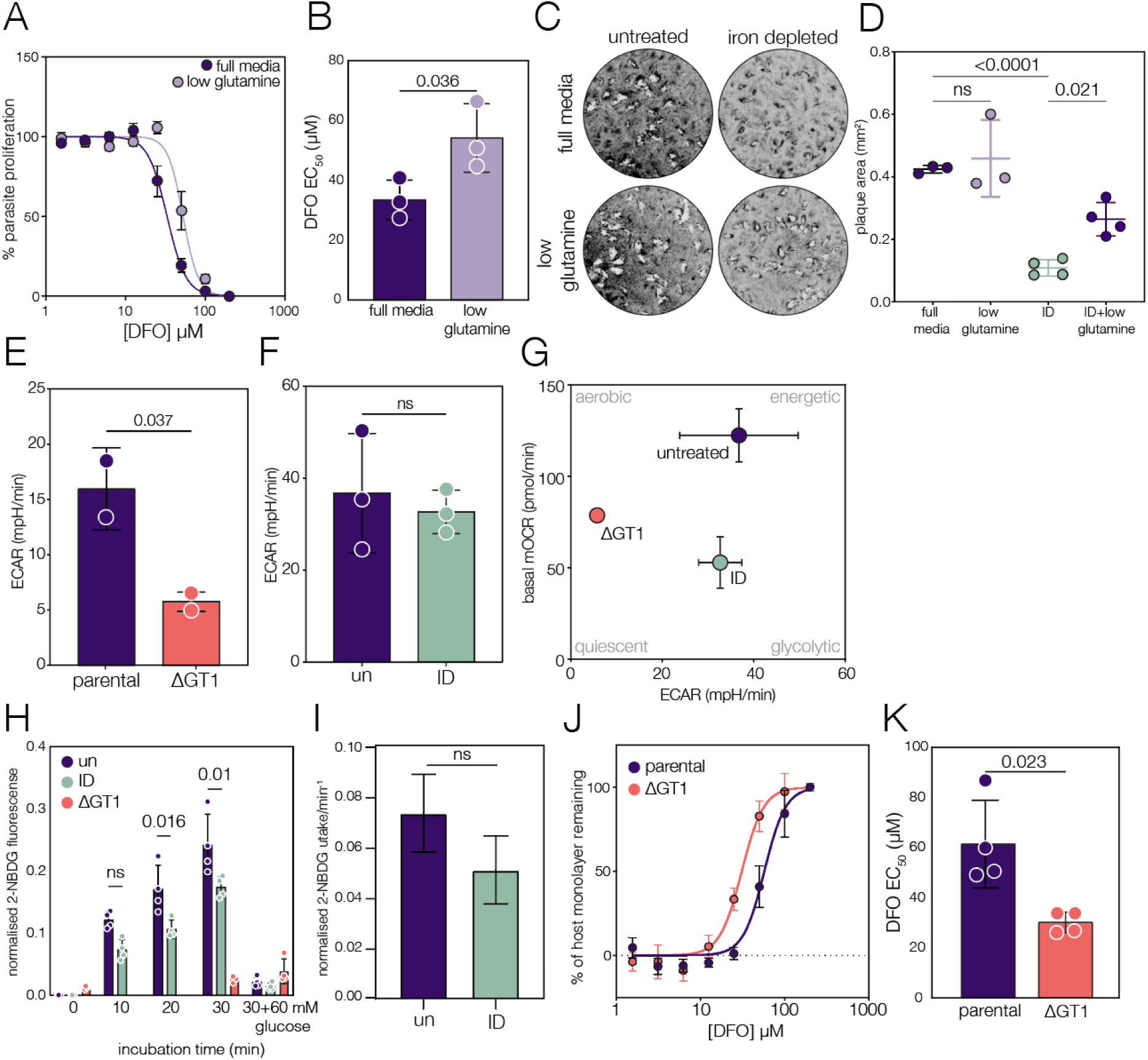
Glucose but not glutamine is required for survival under low iron. (**A**) Fluorescent growth assays to quantify sensitivity to DFO. Mean parasite proliferation in full media (dark) or low glutamine media (light), N=X, ±SEM. (**B**) Bar graph of DFO EC_50,_ each point represents a replicate, bars at mean ±SD. *p* value from two-tailed paired t test. (**C**) Representative plaque assays to assess the growth of ID parasites in different glutamine conditions. (**D**) Quantification of plaque area, bars at mean±SD, *p* values from Brown-Forsythe and Welch ANOVA. No difference observed between full media and low glutamine groups under iron replete conditions (N=3). Co-limitation of glutamine and iron significantly increased plaque area compared to ID alone (N=4). (**E**) Extracellular acidification rate (ECAR) measured by Seahorse assay for parental and ΔGT1 parasites, bars at mean±SD, *p* value from two-tailed t test (N=2). (**F**) ECAR assessed in untreated and iron depleted (ID) parasites, no significant difference by two-tailed t test (N=3). (**G**) Metabolic map with points representing the mean basal mitochondrial oxygen consumption rate (mOCR) on the y-axis and mean ECAR on the x-axis, ±SD, N=3. (**H**) Geometric mean parasite fluorescence upon incubation with fluorescent glucose analog 2-NBDG. Points indicate biological replicates, bars at mean ±SD, *p* values from two way ANOVA with Šídák’s multiple comparisons correction. (**I**) Mean hill slope of 2-NBDG uptake, N=4 (**J**) Quantification of monolayer disruption after infection upon DFO treatment. Points at mean±SEM and data were fitted with a non-linear regression model (N=4). (**K**) Mean DFO EC_50_ for parental and ΔGT1 parasites. Bars represent ±SD, *p* values from two-tailed paired t test (N=4).

The metabolism of glutamine results in the production of ammonia as a by-product of the conversion to a-ketoglutarate (Cheng, Kelsey and Guo, 2022). To investigate this further, we replaced glutamine with the synthetic dipeptide alanyl-glutamine (GlutaMAX) which breaks down more slowly, preventing ammonia buildup but providing increased alanine to the cells. In this case, modulation of alanyl-glutamine did not sensitise cells to iron restriction (**Fig. S4B**). The difference between glutamine and GlutaMAX supports our hypothesise that the carbon source has a direct impact on sensitivity to iron restriction.

These data show glutaminolysis is dispensable to iron starved parasites, and suggests that excess glutamine catabolism in iron deprived cells is potentially toxic.

### *Toxoplasma* requires glucose metabolism during iron deprivation

Unlike mitochondrial respiration, energy generation through glycolysis—although less efficient—does not require iron. To assess if iron deprived parasites were utilising glycolysis, we quantified the extracellular acidification rate (ECAR) using the Seahorse analyser. This gives an indirect assessment of glycolysis based on media acidification after lactic acid excretion and can be used to assess shifts in glucose metabolism (Hayward *et al*., 2022). To confirm that ECAR is an effective proxy for glycolysis, we generated a parasite line which lacks the major glucose transporter, GT1 (TGGT1_214320) (**Fig. S4C**). Previous metabolic studies have shown that these parasites are significantly inhibited in glucose uptake and consequently have a very low rate of glycolysis (Blume *et al*., 2009; Nitzsche *et al*., 2016). ΔGT1 parasites had a small (t test, *p* = 0.031) defect in basal oxygen consumption rate (**Fig. S4D and E**). To ensure the mitochondrial electron transport chain was functional, we tested for complex IV activity (**Fig. S4F**) and ATPsynthase complex integrity (**Fig. S4G**) and saw no changes. However, loss of GT1 led to a significant drop (t test, *p* = 0.037) in extracellular acidification rate (**Fig. 6E**), validating ECAR as a proxy for glycolysis in *Toxoplasma*. In contrast, although iron deprived parasites had a significant defect in mitochondrial respiration, extracellular acidification was maintained (**Fig. 6F**), pushing cells towards a more glycolytic phenotype, as could be seen from the metabolic map (**Fig. 6G**).

Glycolysis requires glucose uptake, to quantify glucose uptake, we made use of the fluorescent glucose analogue 2-NBDG (Oh and Matsuoka, 2002). To validate this approach, we confirmed that ΔGT1 parasites were unable to take up 2-NBDG, and that 2-NBDG uptake could be entirely blocked by excess (60 mM) unlabelled glucose (**Fig. 6H**). Both untreated and iron deprived parasites took up 2-NBDG, although we saw significantly (two way ANOVA with Sidaks correction) less signal at both 20 and 30 mins post incubation in iron deprived parasites (**Fig. 6H**). However, calculating the rate of 2-NBDG uptake showed no significant change in 2-NBDG uptake between conditions (**Fig. 6I and S4E**), demonstrating that iron deprived parasites can effectively take up glucose.

In *Toxoplasma*, although the glycolysis pathway has been maintained, it is typically dispensable under standard growth conditions (Blume *et al*., 2009). To determine if glycolysis becomes more fitness conferring under low iron, we tested the sensitivity of parasite to iron chelation under media containing high (4.5 mM) or low (1 mM) glucose. We saw a small but significant (*p* = 0.0005, extra sum-of-squares F test) increase in sensitivity to iron chelation in low glucose media (**Fig. S4F**). However, changing extracellular glucose concentrations may not directly impact the parasite, and so we made use of the ΔGT1 parasite line. We find that loss of the glucose transporter makes the parasites significantly (*p* <0.0001, extra-sum-of-squares F test) more sensitive to iron chelation (IC_50_ diff. = -31.24 µM) than the parental parasite line (**Fig. 6J and K**), demonstrating that glycolysis is required for replication under low iron conditions.

These results demonstrate the importance of central carbon metabolism in parasite replication under low iron, and highlights the interconnections between metabolism and iron utilisation of this parasite.

## Discussion

Here, we have mapped the effects of acute iron deprivation on the proteome and metabolism of *Toxoplasma gondii.* We find that acute iron deprivation leads to significant remodelling of the parasite proteome, and the changes that we see are poorly correlated with changes in the parasite transcriptome. Poor correlation between protein and transcript levels have previously been observed in other eukaryotes, (Cheng *et al*., 2016; Teh *et al*., 2025), especially upon stress when changes in mRNA availability and translation can lead to rapid changes in the proteome (Williams and Rousseau, 2024).

One potential mechanism for this discordance upon iron deprivation is the rapid and profound drop in parasite translation upon iron deprivation. Translational inhibition upon iron deprivation has been observed in other organisms (Romero *et al*., 2020; Barlit *et al*., 2024; Puig-Segui *et al*., 2024), however the mechanisms and pathways vary. In mammalian cells, translation is inhibited by iron deprivation via signalling through the mTOR pathway (Puig-Segui et al., 2024), which is absent from *Toxoplasma*. In yeast, iron-mediated translation-inhibition is initiated by repression of Rli1 (an ABCE1 homolog) through RNA-binding proteins (Barlit et al., 2024). In *Toxoplasma*, we find that ABCE1 is significantly downregulated upon iron depletion, however this decrease in abundance is seen after repression in translation. Although ABCE1 can be regulated beyond abundance (Barthelme *et al*., 2011; Wu *et al*., 2018), this suggests that translational inhibition precedes ABCE1 changes. However, due to the involvement of numerous iron-containing proteins in regulating translation, including ABCE1, we cannot yet be certain of the mechanism linking iron and translation inhibition. In mammalian cells, the integrated stress response (ISR) also has an important role in the response to low iron (Wang *et al*., 2022) and a number of previous studies have found evidence for an equivalent ISR in *Toxoplasma* (Sullivan *et al*., 2004; Konrad, Wek and Sullivan, 2011, 2014; Augusto *et al*., 2021). This makes the ISR an attractive candidate for mediating iron-dependent translational inhibition. Future work will seek to integrate how the parasite senses iron, and how translation is regulated in response.

A key destination for iron within the parasite is the mitochondrion, where iron is required in enzymes of the TCA cycle and the ETC (Usey and Huet, 2022; MacLean *et al*., 2025; Sloan *et al*., 2025). Here, we find that *Toxoplasma* iron deprivation leads to dynamic and reversible changes in parasite mitochondrial morphology, membrane potential, and in respiration. While crucial for oxidative phosphorylation, mitochondrial polarisation is also vital for the import of proteins, ions and metabolites (Kulawiak *et al*., 2013; Zorova *et al*., 2018) and the depolarisation may be linked to the metabolic phenotypes observed in this study. We also see a drop in both basal and maximal oxygen consumption, which is also seen upon depletion of the major iron and zinc importer ZFT (Aghabi *et al*., 2025), highlighting the importance of iron to mitochondrial respiration.

The changes seen in mitochondrial respiration are reflected in the significant alteration in parasite metabolism seen in metabolomics. We see significant changes in the TCA cycle, including accumulation of citrate and fumarate. Citrate is produced in the mitochondrion by citrate synthase 1 and 2-methylcitrate synthase, however the dispensability of these enzymes (Lyu *et al*., 2024) hints at additional mechanisms of mitochondrial citrate supply. While we cannot say definitively where citrate accumulates in iron depleted cells, citrate is converted isocitrate through the action of aconitase which, although has no change in abundance upon iron depletion, does demonstrate decreased activity (Sloan *et al*., 2025) and increased sensitivity to chemical inhibition.

Citrate also plays a key role in the cytosol where it is used to generate acetyl-CoA through the action of ATP-citrate lyase (ACL), an important step in fatty acid metabolism (Tymoshenko *et al*., 2015; Williams and O’Neill, 2018). Citrate accumulation has previously been associated with an increase in lipid droplets in iron depleted mammalian cells (Crooks *et al*., 2018; De Bortoli *et al*., 2018) and here we report this association in *Toxoplasma*. We see downregulation of ACL at the protein level which could also contribute to an additional accumulation of cytosolic citrate. Alongside downregulation of ACL, we see upregulation of acetyl CoA synthase (ACS) which provides an alternate supply for cytosolic acetyl CoA as ACS utilises acetate scavenged from the host to supply fatty acid synthesis (Dubois et al., 2018). ACS and ACL form an synthetically lethal pair (Tymoshenko et al., 2015), therefore their divergent regulation at the protein level suggests that scavenged acetate is the major driver of fatty acid metabolism and lipid accumulation in iron deprived parasites (Renaud *et al*., 2024).

We also see accumulation of fumarate and decreased abundance of the mitochondrial Fe-S protein SDHB, as has previously been observed in mammalian macrophages (Pereira *et al*., 2019). SDHB abundance has a central role in connecting cellular metabolism between the TCA cycle and the ETC, and is also reduced upon knockdown of the *Toxoplasma* iron and zinc transporter, ZFT (Aghabi *et al*., 2025). SDHB acquires its Fe-S from the mitochondrial (ISC) Fe-S cluster biogenesis (Pamukcu *et al*., 2021) pathway which suggests that both the cytosolic and mitochondrial Fe-S pathways are impacted by iron deprivation.

Beyond mitochondrial metabolism, we also see changes in glycolysis and the pentose phosphate pathway (PPP), including upregulation of the enzyme G6PDH2 and accumulation of several PPP metabolic intermediates. This may be due to the lack of utilisation of nucleotides as DNA replication is blocked or alternatively may have a link to the oxidative state of the parasite. Under standard growth conditions, *Toxoplasma* balances glycolysis and glutamine catabolism to generate energy for invasion and replication (MacRae *et al*., 2012; Nitzsche *et al*., 2016). We find that while mitochondrial respiration is inhibited in the absence of iron, parasites switch to glycolysis for energy generation. Highlighting its importance, parasites unable to take up glucose (ΔGT1) are significantly more susceptible to iron depletion while glutamine restriction, which is predicted to increase flux through glycolysis (Nitzsche *et al*., 2016; Uboldi *et al*., 2025), is protective against low iron. Recently, it was shown that genetic manipulation of glycolysis inhibited virulence in a mouse model, even though growth *in vitro* was not affected (Zhang *et al*., 2025). Iron is frequently withheld during infection (Frost and Drakesmith, 2025), and potentially limited iron during *in vivo* infection could help explain some of the disparate phenotypes seen. This highlights the importance of metabolic flexibility to Toxoplasma, providing a hypothesis for the parasite’s maintenance of these overlapping and frequently redundant pathways.

Here, we show for the first time the consequences of acute iron deprivation on *Toxoplasma*. Iron deprivation leads to a rapid inhibition of parasite translation, proteomic remodelling and a metabolic switch. The battle for iron is an essential aspect of pathogenesis and survival in iron limited environments important for a range of pathogens, including *Toxoplasma*. Future studies will focus on how iron is sensed and the upstream signalling processes required to trigger the large-scale changes that we see here.

## Materials and methods

### *T. gondii* and host cell maintenance

*Toxoplasma gondii* tachyzoites were maintained at 37 °C with 5% CO_2_ and were grown in human foreskin fibroblasts (HFFs) cultured in Dulbecco’s modified Eagle’s medium (DMEM), supplemented with 3% (D3) heat-inactivated foetal bovine serum (FBS), 4 mM L-glutamine, 100 units / ml penicillin and 100 µg/ml streptomycin. HFFs were passaged in DMEM with 10% FBS (D10), 4 mM glutamine and penicillin/streptomycin. For iron deprivation (ID), confluent host cells were incubated with D3 media containing 100 µM deferoxamine (Deferoxamine mesylate salt, Sigma) for 24 hours prior to infection. Iron complemented host cells (ID+FAC) were incubated with both 100 µM deferoxamine and 200 µM ferric ammonium citrate (FAC). For low glutamine conditions, D3 media was supplemented with 0.4 mM L-glutamine.

### Strain construction

To generate gt1 knockout strain, a gRNA targeting the N-terminus of TGGT1_214320 was identified using ChopChop (https://chopchop.cbu.uib.no/) and cloned into the Tub-Cas9YFP-pU6-ccdB-tracrRNA plasmid (Curt-Varesano *et al*., 2016) via BSaI restriction site using Primers P1 and P2. (sgRNA sequences are listed in **Table 3**).

**Table 3.**
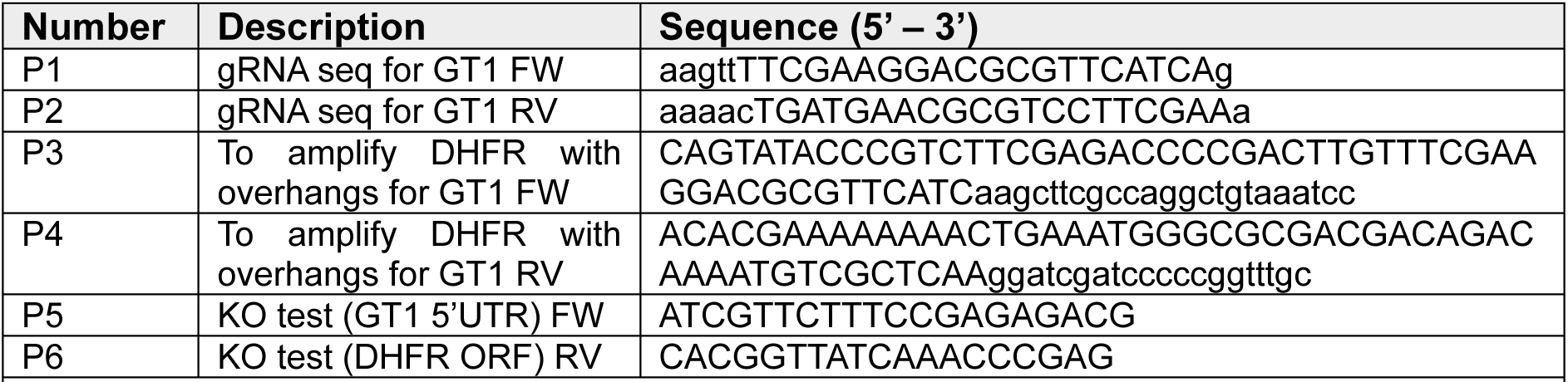
List of gRNA and primers used in this study.

The repair template containing **DHFR** was amplified by PCR with primers P3 and P4 (**Table 3**) containing overhangs encoding 50 bp of homology for the regions upstream and downstream of the TGGT1_214320 ORF. The ΔGT1 strain was generated in a TATi/ΔKu80 background (Sheiner *et al*., 2011) by transfecting with 50 µg of gRNA plasmid generated above and the PCR amplified repair template containing DHFR ORF. Transfections were carried out using a Gene Pulse BioRad electroporator. Parasites were selected using 2 µM pyrimethamine and cloned by serial dilution. Positive clones were tested by PCR using primers P5 and P6 (**Table 3**).

### Immunoblotting

For western blot analysis, parasites were grown on untreated, iron deprived or iron complemented host cells for 24 h. Parasites were mechanically released by scraping and passing 2-3x through a 25 g needle, filtered through a 3 µm polycarbonate filter to remove host cell debris and washed. For puromycin incorporation, 5 x 10^6^ parasites were incubated in media containing 10 µg/ml puromycin (P8833, Sigma) for 30 min at 37 °C prior to lysate collection. Whole cell lysates were prepared by incubating in lysis buffer for 30 mins on ice (150 mM sodium chloride, 1% Triton X-100, 0.5% sodium deoxycholate, 0.1% sodium dodecyl sulphate (SDS) and 50 mM Tris, pH 8.0). An appropriate volume of 4X SDS loading dye was added and samples were boiled for 10 mins. Proteins were separated on a 10% SDS gel before transferred to nitrocellulose membrane for 1 h, blocking at for 1 h at room temperature in blocking buffer (recipe). Membranes were incubated overnight with anti-puromycin (1:1000, MABE343, Merck) anti-*Tg*CDPK1 (1:10,000, (Waldman *et al*., 2020)), anti-HA (1:1000, 11867423001, Merck) or anti-Flag (1:1000, MA191878, Invitrogen) antibodies in blocking buffer and washed three times in PBST. Fluorescent secondary antibodies (IRDye800 and IRDye680, LI-COR) were used to visualise proteins using the Odyssey CLx. For puromycin incorporation, total protein was visualised using ponceau, prior to probing, and quantified by densitometry analysis using ImageJ (Fiji) and used to normalise puromycin signal. Quantification for proteins of interest were normalised to respective loading controls and each condition was plotted relative to the sum of normalised protein quantified in each replicate to show variation in controls between replicates, as previously described (Degasperi *et al*., 2014).

Complex-V (ATP-Synthase) assembly was detected using Blue-Native PAGE followed by western blot. Briefly, freshly extracellular parasites were solubilised in 1% β-DDM (w/v) in 750 mM aminocaproic acid solution on ice for 10 minutes. The samples were centrifuged at 16,000 × g at 4°C for 30 minutes in a bench-top centrifuge. The resulting supernatants were combined with Coomassie G250 to a final concentration of 0.25%, and proteins were resolved on pre-cast 4–16% NativePAGE™ Bis-Tris gels (Invitrogen). The gel was run at 80V for 1 hour followed by 250V for 2 hours. NativeMark™ Unstained Protein Standard (Invitrogen) was used as a molecular weight marker. Proteins were transferred to a 0.45 μm PVDF membrane (Hybond, Merck) using the wet transfer method at 100 V for 1 hour in Towbin transfer buffer (25 mM Tris, 192 mM glycine, 10% methanol). Membranes were blocked using 5% (w/v) skimmed milk in PBS with 0.1% Tween-20, probed with a primary rabbit anti-ATP-β antibody (1:2000, Agrisera AS05085), followed by a horseradish peroxidase (HRP)-conjugated secondary anti-rabbit antibody (1:10,000, Promega W4011). Detection was done using the Pierce ECL substrate (Thermo Scientific).

### OPP incorporation

ΔKu80 parasites were grown on coverslips with untreated, iron deprived or iron complemented host cells for 24 h. Cells were then labelled with 20 µM Click-iT O-propargyl-puromycin conjugated to Alexa Fluor 488 (OPP, Invitrogen) for 30 min at 37 °C. Cyclohexamide (CHX) was used as a negative control, with parasites incubated with 100 µM CHX for 20 min before OPP labelling. Coverslips were washed with PBS before being fixed with 4% formaldehyde and permeabilised with 0.5% Triton X-100. Using a click chemistry reaction, OPP was labelled following manufactures instructions with Alexa Fluor® picolyl azide for 30 min at room temperature, protected from light. Coverslips were mounted with Fluoromount with DAPI (Southern Biotech) and imaged using a Zeiss LSM 880 inverted confocal microscope and Zen black software (Zeiss). Images were processed and analysed with an automated macro with ImageJ (Fiji).

For the OPP incorporation and ABCE1 time course experiment, host cells on coverslips were left untreated or pretreated for 24 hours prior to infection with ABCE1-HA parasites. After 24 hours, iron depleted medium was removed and wells were washed with fresh media twice before fresh media was added to begin the recovery process. At 6, 12, 18 and 24 hours after infection or recovery, coverslips were labelled with OPP as described in the previous paragraph. Coverslips were then incubated 0.5% BSA in PBS containing anti-HA (1:1000, Merck 11867423001) and anti-TgIMC1/TgSAG1 (1:1000) antibodies. Coverslips were then washed with PBS before incubation with anti-rat and anti-mouse secondary antibodies conjugated to Alexa Fluor 594 or Alexa Fluor 647 respectively (both 1:1000). After further PBS washes, cells were then mounted onto slides with Fluoromount with DAPI (Southern Biotech). Images were obtained using the Zeiss LSM 880 inverted confocal microscope using Zen black software (Zeiss). Images were obtained from 4 replicates with each replicate consisting of the vacuoles contained within 5 images. OPP incorporation and HA abundance were quantified in parasite vacuoles using automated macros within ImageJ (Fiji). For HA abundance, HA signal contained within host cell nuclei was used to background adjust the signal obtained from parasite within the same image.

### Seahorse extracellular flux analysis

Mitochondrial oxygen consumption rate (mOCR) and extracellular acidification rate (ECAR) were measured with a Seahorse XF HS Mini analyser (Agilent) as previously described (Hayward *et al*., 2022; Silva *et al*., 2023). Briefly, parasites were grown as indicated for 24 hours before being released and filtered as above and washed with Seahorse XF DMEM base medium supplemented with 5 mM glucose and 1 mM glutamine. A total of 1.5 x 10^6^ parasites were added to each well of an 8-well XF miniplate pretreated with poly-L-lysine and OCR and ECAR quantified. The background, non-mitochondrial oxygen consumption rate was defined as that present after addition of 1 µM atovaquone. Basal and maximal mOCR were calculated by taking the OCR measurement before (basal) or after (maximal) addition of 1 μM FCCP and subtracting the background rate. Each experiment was performed in triplicate, with 3 biological repeats.

### Fluorescence growth assays

HFF cells were seeded onto clear-bottom black 96-well plates (Thermo Scientific), allowed to reach confluency and infected with 1000 mNeon or tdTomato (Aghabi *et al*., 2023) parasites per well. After 2 h at 37 °C, the media was replaced with indicated concentrations of drug added. After 4 days of infection, parasite fluorescence was measured using a PHERAstar FS microplate reader (BMG LabTech) at 488 or 594 nm emission for mNeon and tdTomato respectively. Experiments were performed in technical triplicate with at least 3 biological replicates. Uninfected wells served as blanks to remove background fluorescence and values were normalised to infected, untreated wells. Dose response curves and EC_50_ calculations were performed in GraphPad Prism 10, using the inhibitor vs. normalised response (variable slope) model. Extra sum of squares F-tests or t tests were performed to test differences in best-fit IC_50_ values between groups.

### Flow cytometry analysis

Glucose uptake was assessed by flow cytometry using the fluorescent glucose analogue 2-NBDG (Invitrogen, N13195). ΔKu80::tdTomato or ΔGT1 parasites were grown on untreated or iron depleted host cells for 24 hours. Parasites were released by scraping and syringing infected host cells and parasites were filter from host material through a 3 µm filter. Parasites were washed in glucose free media before being resuspended in DMEM supplemented with 2.5 µM l-glutamine only. As a competitive inhibition control, parasites were resuspended in 2.5 µM l-glutamine and 60 mM glucose. Parasites were then left unstained or were incubated with 1 mM 2-NBDG for 10, 20 or 30 minutes at 37 °C. Competition controls incubated with 1 mM 2-NBDG for 30 minutes. Parasites were analysed on a BD FACSCelesta Flow Cytometer and data was acquired using FACSDiva software (BD Biosciences). Parasites were gated on forward and side scatter and on red fluorescence (only for tdTomato parasites). All data were analysed using FlowJo v10 (BD Biosciences) and the geometric mean of the BB515 channel was used to quantify 2-NBDG uptake.

### Immunofluorescence assay

Immunofluorescence assays were performed on both intracellular and egressed extracellular tachyzoites. Intracellular *T. gondii* parasites were grown on coverslips pre-seeded with HFFs before being fixed with 4% paraformaldehyde for 10 minutes at room temperature. Cells were permeabilised for 10 mins in 0.5% Triton X-100 in PBS before being stored in 2% BSA, 0.2% Triton X-100 at 4 °C overnight. Coverslips were then incubated with primary antibodies in PBS; rat anti-HA (1:1000, Merck 11867423001), guinea pig anti-*Tg*CDPK1 (1:5,000), mouse anti-*Tg*IMC1/*Tg*SAG1 (1:1000) or rabbit anti-*Tg*TOM40 (1:1000). Coverslips were then washed with PBS before incubation with secondary antibodies raised against the host of the corresponding primary antibody, conjugated to Alexa Fluor 488, Alexa Fluor 594 or Alexa Fluor 647 (all 1:1000). After further PBS washes, cells were then mounted onto slides with Fluoromount with DAPI (Southern Biotech). Images were obtained using the Zeiss LSM 880 inverted confocal microscope using Zen black software (Zeiss) or with Leica DiM8 (Leica Microsystems) microscope and processed using LasX (Leica Microsystems). Images were processed and analysed with an automated macro with ImageJ (Fiji).

For time course analysis of mitochondrial morphology, host cells on coverslips were left untreated or pretreated for 24 hours prior to infection with ΔKu80 parasites. After 24 hours, iron depleted medium was removed from recovery wells, cells were washed with fresh media twice before fresh media was added to begin the recovery process. At 6, 12, 18 and 24 hours after infection or recovery, cells were fixed, permeabilised as described in the previous paragraph. Coverslips were stained with anti-TOM40 (1:1000) and anti-CDPK1 (1:5000) antibodies before washing in PBS before incubation with secondary antibodies.

### MitoTracker Imaging

Parasites treated as above were mechanically released, washed once with PBS and incubated in the presence of 10 µM Hoechst 33342 (ThermoFisher Scientific, H3570) and 50 nM MitoTracker Deep Red (Invitrogen, M22426) for 10 mins at 37 °C on poly-L-lysine treated glass bottom dishes. For imaging of host mitochondrial association, ΔKu80:tdTomato parasites were cultured for 24 hours on untreated, ID or ID+FAC host cells before incubation with 50 µM MitoTracker Green FM for 15 mins at 37 °C. Parasites were imaged live using Leica DiM8 (Leica Microsystems) microscope and processed using LasX (Leica Microsystems). Intensity was analysed manually using ImageJ (Fiji).

### Crystal violet growth assays

HFF cells were seeded onto clear 96-well plates (Thermo Scientific), allowed to reach confluency and infected with 1000 ΔGT1 or parental TATi/ΔKu80 parasites per well. After 2 h at 37 °C, the media was replaced with indicated concentrations of drug added. After 4 days of infection, media was removed and cells were fixed in 20 µl ice-cold methanol before being stained for at least 2 hours in 0.5% crystal violet in 20% ethanol solution. Wells were then washed with sterile water before being left to dry. Absorbance was measured in a PHERAstar FS microplate reader (BMG LabTech). Experiments were performed with 6 technical repeats with 4 biological replicates. Unstained wells served as blanks to remove background and values were normalised to infected wells treated at 200 µM DFO. Dose response curves and IC_50_ calculations were performed in GraphPad Prism 10, using the inhibitor vs. normalised response (variable slope) model. Extra sum of squares F-tests were performed to test differences in best-fit IC_50_ values between groups and differences in average IC_50_ were analysed by t-test.

### Plaque assays

1000, 500 or 250 parasites were dispensed onto confluent monolayers of HFF cells in 6-well plates. Parasites were allowed to replicate for 5-7 days undisturbed before being washed with PBS followed by fixation with ice cold methanol for 10 minutes. Cells were stained using 0.5% crystal violet in 20% ethanol solution for at least 2 hours at room temperature before washing with distilled H_2_O. Plates were imaged and plaque numbers and sizes were quantified using ImageJ (Fiji).

### Proteomics sample preparation

To prepare samples for proteomics 1 x 15 cm dishes for control conditions and 2 x 15 cm dishes for iron deprived conditions were pooled to generate each replicate. For iron deprivation, confluent host cells in 15 cm dishes were incubated with D3 media containing 100 µM deferoxamine (Deferoxamine mesylate salt, Sigma) for 24 hours prior to infection. After 24 hours of infection, cells were washed twice in PBS to remove extracellular parasites, before being scraped and passed through a 25 gauge needle to mechanically release parasites. Host cell material was filtered from parasites by passing through a 3 µm filter. Parasites were collected by centrifugation at 3000 xg for 25 minutes. Parasites were counted and 4 x 10^7^ parasites were resuspended in lysis buffer (50 mM K phosphate buffer, 1% SDS, 1 mM EDTA, 1 mM DTT, 1 x Universal nuclease (Thermo Scientific, 12156490), 1 x Halt protease inhibitors (Thermo Scientific, 10720825) and left to incubate on ice for 30 minutes. Lysates were freeze thawed three times in a dry-ice and ethanol bath to enhance lysis.

Insoluble material was removed from lysates by centrifuging at 20,000 xg at 4°C for 20 minutes and collecting the supernatant. Samples were then reduced in 25 mM TCEP (Fisher) for 10 minutes at 37°C and then alkylated with 25 mM iodoacetamide (Fisher) for 1 hr at RT in darkness. Proteins were then precipitated by adding 12% volume ice-cold trichloroacetic acid (TCA) (Sigma) and incubated at -20°C overnight. Precipitated proteins were pelleted at 16,000 xg at 4 °C washed with agitation 3 times with ice-cold acetone. Samples were then resuspended in 100 mM triethylammonium bicarbonate (TEAB) (Fisher) and were then digested by a Trypsin/Lys-C mix (Promega) at a 1:25 (w/w) ratio overnight at 37°C. Peptides were dried using a SpeedVac SPD1030 vacuum concentrator. Peptide samples were resuspended in 1% formic acid (#85178, Thermo Scientific) before being subjected to quality control.

### Complex IV activity assay

Mitochondrial complex-IV activity was assessed using an in-gel enzymatic assay as previously described (Lacombe *et al*., 2019). Briefly, freshly extracellular parasites were solubilised in a buffer containing 50 mM NaCl, 2 mM 6-aminohexanoic acid, 50 mM imidazole, 2% (w/v) β-DDM, and 1 mM EDTA–HCl (pH 7.0), incubating on ice for 10 minutes. After incubation, the samples were centrifuged at 16,000 × g for 15 minutes at 4°C in a bench-top centrifuge. The supernatants were transferred to new tubes and mixed with a loading buffer containing 6.25% glycerol and 0.125% Ponceau S. Samples were loaded onto pre-cast 4-16% NativePAGE™ Bis-Tris gels (Invitrogen) and run for 1 hour at 100V, followed by 250V until the dye front ran out of the gel (∼1.5hrs). To detect complex-IV activity, the gel was incubated in a freshly prepared and pre-warmed (37°C) staining solution containing 50 mM potassium phosphate buffer (pH 7.2), 1 mg/ml cytochrome c, and 0.1% (w/v) 3,3′-diaminobenzidine tetrahydrochloride (DAB) for ∼3 hours before imaging.

### Mass Spectrometry

Analysis of the peptides was performed on a Q Exactive HF Quadripole Orbitrap Mass Spectrometer (Thermo Scientific) coupled to a Dionex Ultimate 3000 RS HPLC system (Thermo Scientific). The buffers used were Buffer A (0.1% formic acid in Milli-Q water (v/v)) and Buffer B (80% acetonitrile (#83640.290, VWR) and 0.1% formic acid in Milli-Q water (v/v). An equivalent of 1.25 µg of peptides from each sample were loaded at 10 µl/minute onto an Acclaim PepMap 100 C18 column (#164750, Thermo Scientific). This trapping column was washed for 6 minutes at the same flow rate with 1% Buffer B and then switched in-line with a µPAC Neo column (#COL-NANO110NEOB, Thermo Scientific). Buffer B was set to 3.8% and then increased over the next 117 minutes to 41.3%; increased to 61.3%, 99% in 13 minute and 10 minute steps; the column was then washed for 17 minutes. Two blanks were run between each sample to reduce carry-over. The column was kept at a constant temperature of 50 °C.

The Q-Exactive HF was operated in positive ionisation mode using easy spray source. The source voltage was set at 1.90 kV and the capillary temperature was 250 °C. Data were acquired in Data Independent Acquisition Mode as previously described (Doellinger et al., 2020), with little modification. A scan cycle comprised a full MS scan (345 – 1155 m/z), resolution was set to 60,000, AGC target 3 x 106, maximum injection time 200 ms. MS survey scans were followed by DIA scans of dynamic window widths with an overlap of 0.5 Th. DIA spectra were recorded at a resolution of 30,000 at 200 m/z, using an automatic gain control target of 3 x 106, a maximum injection time of 55 ms and a first fixed mass of 200 m/z. Normalised collision energy was set to 25% with a default charge state set at 3. Data for both MS scan and MS/MS DIA scan events were acquired in profile mode.

### Proteomics data analysis

Analysis of the DIA data was carried out using Spectronaut (version 17.4.230317.55965, Biognosys, AG). The directDIA workflow, using the default settings (BGS Factory Settings) with the following modifications was used: decoy generation set to inverse, Protein LFQ Method was set to QUANT 2.0 (SN Standard) and Precursor Filtering set to Identified (Qvalue); Precursor Qvalue Cutoff and Protein Qvalue Cutoff (Experimental) set to 0.01; Precursor PEP Cutoff set to 0.1, Protein Qvalue Cutoff (Run) set to 0.05 and Protein PEP Cutoff set to 0.75.

For the Pulsar search the settings were: maximum of 2 missed trypsin cleavages; PSM, Protein and Peptide FDR levels set to 0.01; scanning ranges et to 300-1800 m/z and Relative Intensity (Minimum) set to 5%; cysteine carbamidomethylation set as fixed modification and acetyle (N-term), deamidation (asparagine, glutamine), dioxidation (methionine, tryptophan), glutamine to pyro-Glu and oxidation of methionine set as variable modifications. The databases used were T.gondiiGT1 (reviewed, downloaded from ToxoDB on 2023-09-14).

Differential expression analysis was performed in RStudio using the ProteoDA package (Thurman *et al*., 2023). Protein intensity values were evaluated using proteiNorm tool (Graw *et al*., 2020) before being transformed using variance stabilising normalisation. Statistical analysis was performed using the limma package (Ritchie *et al*., 2015) in R. Proteins with an FDR adjusted *p*-value <0.05 and a fold change >1.5 (log_2_0.6) were considered differentially expressed. DISCO scores were calculated by integrating these proteomic data with transcriptomic data previously reported in (Sloan *et al*., 2025). DISCO scores were computed as previously described (Domaszewska *et al*., 2017) using the formula DISCO score = log_2_FCRNA x log_2_FCPROT x (log_10_p-adjRNA + log_10_p-adjPROT) x sign(log_2_FCRNA) x sign(log_2_FCPROT).

### Metabolomics sample preparation

For metabolite extraction, 2 x 15 cm dishes for control conditions and 4 x 15 cm dishes for iron deprived conditions were pooled to generate each replicate. For iron deprivation, confluent host cells in 15 cm dishes were incubated with D3 media containing 100 µM deferoxamine (Deferoxamine mesylate salt, Sigma) for 24 hours prior to infection. After 24 hours of infection dishes were moved to a shallow container of ice water to quench metabolism. Cells were washed in ice-cold PBS twice to remove extracellular parasites and old media, before being scraped and passed through a 25 gauge needle to mechanically release parasites. Parasites were filtered from host material through a 3 µm filter before centrifugation at 1 °C for 25 mins at 3000 xg. Pellets were resuspended and washed in 1 ml ice-cold PBS before being counted using a haemocytometer. 4 x 10^7^ parasites were resuspended in 200 µl of a chloroform /methanol/water mixture (1:3:1 ratio). Samples were shaken for 1 hour at 4 °C and centrifuged at 20,000 xg for 5 minutes at 4°C. Metabolites in the supernatant were then collected.

### Metabolomics data analysis

Metabolites were quantified by LCMS as previously described (Pountain and Barrett, 2019). In brief, samples were separated by high performance liquid chromatography on a Dionex UltiMate 3000 RSLC system (Thermo) using a ZIC-pHILIC column (Merck). Mass spectrometry was performed using a Orbitrap Q Exactive (Thermo) with analysis performed in both positive and negative ionisation modes. To improve metabolite identification, samples were run alongside internal standard mixtures that included metabolites from a wide range of metabolic pathways. Additionally, quality control pooled samples were also subject to fragmentation to acquire MS2 spectra using data-dependent acquisition (DDA). Peaks were annotated on the IDEOM platform (Creek *et al*., 2012) and statistical analysis was performed using the MetaboanalystR 4.0 package within RStudio (Pang, Xu, *et al*., 2024). Median normalisation, log10 transformation and mean centering was performed on peak intensities from identified metabolites prior to hierarchical clustering and differential abundance analysis within MetaboanalystR 4.0. Pathway analysis was performed with MetaboAnalyst 6.0 using global test enrichment against the *Toxoplasma gondii* KEGG pathways (Pang, Lu, *et al*., 2024). Relative-betweenness centrality was used as a topology measure of pathway impact. Metabolites closer to each other in the pathway are assigned a higher pathway impact score (between 0 and 1).

### Data availability

The mass spectrometry proteomics data have been deposited to the ProteomeXchange Consortium via the PRIDE partner repository (https://www.ebi.ac.uk/pride/) with the dataset identifier PXD066828. Raw metabolomics MS data are available at the MetaboLights (https://www.ebi.ac.uk/metabolights/) data repository under project MTBLS12831. RNAseq raw data available at EBI ENA (https://www.ebi.ac.uk/ena/browser/home) under project PRJEB67890 and PRJEB83013.

## Acknowledgements

J.H. and C.R.H. are funded by a Sir Henry Dale Fellowship from the Wellcome Trust, the Royal Society (213455/Z/18/Z) and the Carnegie Trust (RIG013355). M.A.S is funded by an Early Career Award from the Wellcome Trust (225677/Z/22/Z). S.S. is funded by the Swiss National Science Foundation Postdoctoral Mobility Fellowship (200158). The funders had no role in study design, data collection and analysis, decision to publish, or preparation of the manuscript.

The authors gratefully acknowledge the MVLS Shared Research Facilities, especially Clement Regnault, for their support and assistance and www.ToxoDB.org without which this work would not have been possible. We thank the Dundee Fingerprints proteomics facility and Michael Hanna for data curation. We thank Sebastian Lourido, Giel van Dooren and David Sibley for providing antibodies and Andrew Maclean and Lilach Sheiner for SDHB-HA, Rieske-HA and ABCE1-HA tagged lines and for helpful discussion and feedback.

## Supplementary figure legends

Figure S1. (**A**) Volcano plot of proteomics data from untreated and iron depleted parasites, highlighting significantly (adj. *p* < 0.05) upregulated (L_2_FC>0.6) (red) and downregulated (L_2_FC<-0.6) (blue) proteins. Each dot represents a protein with larger dots representing iron sulfur cluster (Fe-S) proteins (Renaud, Maupin and Besteiro, 2025) that appear in this dataset. (**B**) Immunofluorescence images showing Gra7 staining (teal) in untreated, iron depleted (ID) and iron complemented parasites (ID+FAC) (red). Gra7 visible inside parasites and on the parasitophorous vacuole, regardless of growth condition. Scale bar 5 µm.

Figure S2. (**A**) Violin plot of mean OPP-fincorporation in untreated or cycloheximide (CHX) treated parasites. Each point represents a parasite vacuole and bars at mean ±SD. *p* value from Welch’s two tailed t-test (*p*<0.0001). (**B**) Plot of normalised (using variance stabilising normalisation (vsn)) ABCE1 protein intensities from MS in untreated and ID parasites. *p* values assigned by fitting a linear model. (**C**) Vacuole area measurements over time in ID and recovering parasites. Each point represents the mean vacuole size/replicate, bars at mean±SD. *p* values from one-way ANOVA with Tukey’s correction. No significant differences were observed between timepoints for ID parasites. Compared to 6 hours of iron depletion, 12, 18 and 24 hours of recovery from iron deprivation coincided with a significant increase in vacuole size (*p*=0.006, *p*=0.0007 and *p*=0.0001 respectively). (**D**) HFF OPP incorporation. Each point represents the mean host cell incorporation from 5 fields of view, line at mean, normalised to parasites at 6 hours. *** indicates significant (*p* < 0.0001) difference between untreated and ID at indicated timepoint. Ns indicates no significant difference between untreated at 6h and recovery at 36 h. *p* values from two way ANOVA with Tukey’s correction. (**E**) Violin plots of OPP in parasite vacuoles. Results from 4 biological replicates for 6, 12, 18 and 24 hours (N=382, 408, 460 and 323). Dotted lines at median and quartiles. (**F**) OPP incorporation and ABCE1 abundance in vacuoles > 20.1 µm^2^. (**G-I**) Fluorescent growth assays testing the efficacy of pyrimethamine (**G**), atovaquone (**H**) or dihydroartemisinin (**I**) on untreated parasites or those recovering from iron depletion. Recovery parasites were previously ID as described, then moved to standard media. (**J**) Bar graph showing EC_50_ of DHA against untreated or recovering parasites. Bars at mean±SD. *p* value from two-tailed paired t test.

Figure S3. (**A**) Fluorescent parasite growth assay assessing the efficacy of aconitase inhibitor sodium fluoroacetate (NaFAc) against parasites cultured in full media and 20 µM DFO (N=4). (**B**) Immunoblot of lysates from Rieske-FLAG parasites, CDPK1 included as a loading control. (**C**) Quantification of Rieske-FLAG, points represent replicates (N=3), bar at mean±SD, *p* value from two-tailed paired t test.

Figure S4. (**A**) Number of plaques counted for each media condition, bar at mean±SD . (**B**) Fluorescent parasite growth assay assessing the efficacy of DFO against parasites cultured in media with 4 mM GlutaMAX and low (0.4 mM) GlutaMAX media (N=3). (**C**) Diagram and PCR confirmation for ΔGT1 strain construction. (**D**) Basal and (**E**) maximal mOCR was determined for both parental and ΔGT1 parasite lines. Each point represents a replicate (N=2) and two-tailed paired t-tests were used to detect significant differences. (**F**) Complex IV assay showing no change in complex IV activity between parental and ΔGT1 parasites. (**G**) Native western blot demonstrating ATPsynthase at correct size for full complex (probed using antibody against ATPsynthase β-subunit. (**H**) Normalised 2-NBDG fluorescence for untreated and iron depleted parasites was plotted for 10, 20 and 30 mins uptake, slope value calculated from linear regression, differenes tested using an extra sum of squares F-test. (**F**) Summarising EC_50_ for DFO in full media and low glucose media. *p* value calculated using a two-tailed t test.

**Table S1** – Summary data from global proteomics from untreated and ID parasites. Predicted protein localisations from (Barylyuk et al., 2020).

**Table S2** – DISCO analysis of correlation between changes in iron depletion between proteomics and RNAseq from (Sloan *et al*., 2025), raw data available at EBI ENA online server under project PRJEB67890 and PRJEB83013.

**Table S3** – Summary data from metabolomics, including raw abundance data (prenorm) and normalised abundance data for identified metabolites. Results from pathway analysis are also included. Raw data are available at the MetaboLights (https://www.ebi.ac.uk/metabolights/) data repository under project MTBLS12831.

